# Demystifying The Myelin *g* ratio: Its Origin, Derivation and Interpretation

**DOI:** 10.1101/2025.02.19.639116

**Authors:** Alexander Gow

**Author notes:** Correspondence: Dr Alexander Gow Center for Molecular Medicine and Genetics, 3216 Scott Hall, 540 E Canfield Ave, Wayne State University School of Medicine, Detroit, MI, 48201.

## Abstract

Most studies involving myelin *g* ratios over the past 120 years assume this metric enumerates changes in myelin thickness (larger *g* ratio = thinner myelin) with axon or fiber diameter. And, moreover, such changes are directly correlated with internodal function (conduction velocity). However, such assumptions are warranted only in the absence of experimental errors and artifacts (i.e. under theoretical conditions). In reality, *g* ratios easily under- or overestimate rates of change exceeding 10%, especially for small caliber fibers. Typical analyses of myelin internodes rely on an explicit mathematical model, 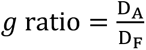, where D_A_ is axon diameter and D_F_ is fiber diameter (myelin plus axon). Shown recently and herein, this model approximates normal physiological conditions only when the axon-fiber diameter relation is directly proportional, whence it is concordant with the axomyelin unit model. However, in transient or non-steady states (development/aging, disease or myelin plasticity) with linear but not directly proportional relations, *g* ratios poorly describe myelin structure. Acceptance of this counterintuitive assertion is predicated on a detailed understanding of the *g* ratio – origins, properties and the biology represented – heretofore uncharted. In light of such *g* ratio limitations, more general and reliable metrics are proposed, the myelin *g_c_* ratio and the *g’* cline.

## Introduction

The myelin *g* ratio represents the gold standard for measuring internodal structure under conditions of steady-state physiology, pathophysiology, development/aging and myelin plasticity in the central and peripheral nervous systems (CNS and PNS, respectively)^1^. It is widely viewed as representing important myelin characteristics, despite a certain *je ne sais quoi*. When myelin biologists use electron microscopy (light microscopy is not recommended^2,3^) to describe the relationship between axons and fibers (axon plus myelin sheath) from *g* ratio plots, they declare a specific mathematical model, 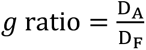, where D_A_ is the axon diameter and D_F_ is the fiber diameter (equivalently, radii or axon-myelin cross-sectional areas). But the simplicity of this equation belies the complexity of several underlying assumptions, which are consequential for both data processing and constrained interpretation of myelin internodal biology^4^.

For example, the *g* ratio plot is only superficially coupled with the physical dimensions of each myelinated axon; rather, it is actually a simple estimator for the slope of the axon-fiber relation (Supplement S2). But there are also deeper connections to the counterposing effects of axon caliber and myelin thickness for a specified fiber diameter, which accounts for innumerable observations that axon-to-myelin ratios fluctuate yet have minimal impact on function^5^. Moreover, the *g* ratio equation (which is a mathematical model) depends crucially on two properties: (1) the axon-fiber diameter relation is directly proportional (i.e. D_A_ ∝ D_F_, where a regression fit to the axon-fiber diameter plot passes through the Origin), and (2) regression fits to *g* ratio plots that are consistent with the first property must be linear with zero slope (i.e. the *g* ratio is invariant and independent of fiber diameter^4^). Fundamentally, if the first property is false, then the mathematical model is false such that *g* ratios derived from the axon-to-fiber diameter relation may be meaningless and cause ambiguous interpretation of the biology.

In a recent study, measurements of myelinated fibers in the optic nerves of adult wild type and *rsh* mice were reexamined from earlier work^6^, to help validate a new analysis pipeline that was developed to minimize statistical artifacts and increase the reliability of *g* ratio interpretation^7^. This methodology relies on a novel model of the myelin internode, designated as the axomyelin unit model^4^, which constrains interpretation of statistical findings associated with myelin internode biology. Under normal physiological conditions and in some disease states (e.g. experimental autoimmune encephalomyelitis assessed in optic nerve), the axomyelin unit model is concordant with experimental measurements. In contrast, measurements in other disease states like the *rsh* mouse model^8^ of the leukodystrophy, Pelizaeus-Merzbacher disease^9^, deviate substantially from axomyelin unit model predictions, suggesting perturbed axon-oligodendrocyte interactions.

These findings notwithstanding, a number of questions have remained unanswered or were brought to light. To answer such questions, the *rsh* dataset used by Gow and colleagues^7^ has been expanded with substantial new data in the analysis performed for the current study. And while those lingering questions are resolved herein, additional statistical and interpretational shortfalls have been uncovered. The deficiencies differ from recently-described artifacts in *g* ratio plots^4^, are widespread in the literature, have persisted for as long as a century and cast doubt on our current understanding about important aspects of myelin internodal biology.

## Methods

### Previous study of rsh data and assurances of humane care of mice

The raw electron microscopy data used in the current study was derived from a previous project^6^ in which the authors stated that all experiments on mice were humane and approved by the Institutional Animal Care and Use Committee at Wayne State University. The axon diameter (D_A_) and myelin radial thickness (doubled to derive the myelin diameter, D_M_) measurements from that study, comprised of medium and large diameter fibers, have been combined with new measurements of small diameter fibers generated for the current study. The legacy data account for 20% of the total data analyzed herein.

### Statistical tests and regression analyses

Raw measurements from digitized electron micrograph negatives (8 nm/pixel resolution) were initially assembled in Microsoft excel (ver 16.66.1) to generate axon and fiber diameter measurements. All further data processing and statistical tests were performed using Graphpad Prism ver. 10.4.1 (Graphpad Software, LLC), including a template for a Data Analysis Pipeline developed previously^7^ which is available on reasonable request. The pipeline uses Deming and nonlinear regression, Welch’s unpaired *t*-tests, mixed-effects analysis with Geisser-Greenhouse epsilon correction and Dunnett’s post hoc tests to assess the results. Deming and nonlinear regression were used for analyzing the axon-fiber diameter relation, mixed-modeling was used for within-group and between-group comparisons to compensate for pseudo-replicates, which is the data structure of all myelin *g* ratio studies. Regression analysis of the skewness in *g* ratio relative frequency histograms was fit using the Gaussian function in Prism, or the Gumbel probability density function coded as^10^:

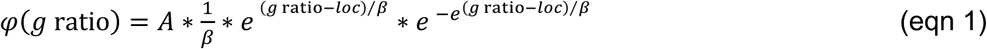

and estimating three parameters: *A*, the function amplitude; *β*, the scale parameter, and; *loc*, the *x*-axis location parameter.

### Box-Cox transformation analysis

A Box-Cox type Lambda transformation series^11^ was used to determine if axon-fiber diameter plots are consistent with a linear relation^4^ and constrained by the biological properties of myelinated fibers. Briefly, axon diameters measured for each mouse are transformed according to the equation:

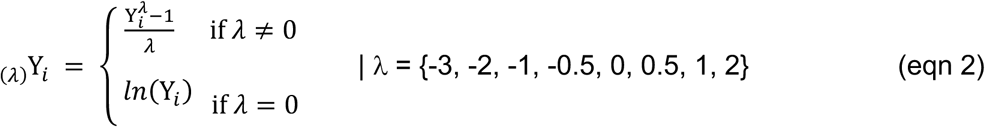

where *λ* = 0 corresponds to ln(Y*_i_*) and _(λ)_Y*_i_* are the transformed axon diameter values. A numerical constant, c, can be added to all Y_)_ (e.g. c = 1), which is convenient for eliminating any occurrences of _(*λ*)_Y*_i_* < 0. Values of _(*λ*)_Y*_i_* are plotted against corresponding fiber diameter values in a summary plot. Pearson’s correlation coefficients and simple linear regression are used to determine *x*- and *y*-intercepts, slopes and *r_xy_* values for each transformation. These parameters are plotted and evaluated for consistency with known biological properties of myelinated fibers.

### Regression fits to g ratio plots

The axomyelin unit model defines the axon-fiber diameter relation as directly proportional under normal physiological conditions^4^. Fiber diameter is represented on the *x*-axis (Supplement S1) so a regression fit to such data can be described by the linear equation:

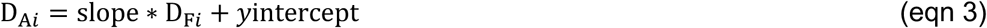

If *y*intercept = 0, the corresponding *g* ratios can be computed as:

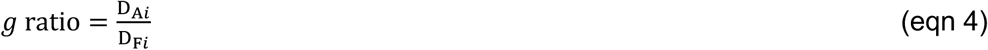

and plotted as a relation of fiber diameter^12,13^. When using this equation, a regression fit to the *g* ratio plot has zero slope, by definition (Supplement S2 and S3).

On the other hand, if the axon-fiber diameter relation is linear but is not directly proportional (e.g. in non-steady states), then the initial regression fit (eqn 3) has a non-zero *y*-intercept (*y*intercept ≠ 0). In such instances, *g* ratios computed using eqn 4 will be markedly asymmetric, and a regression fit will have a non-zero slope or be curvilinear^7^. Such data are appropriately fit using a reciprocal function to account for the *y*-intercept (Supplement S4):

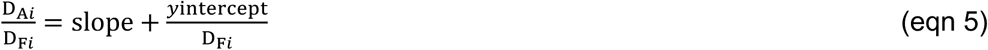

where the slope (from eqn 3) is the maximum value (i.e. the asymptote) of the function as D_F*i*_ increases. Both the slope and *y*-intercept are constants derived from eqn 3.

## Results

To resolve unanswered questions from a recent study involving the pathophysiology of myelinated fibers in *rsh* mice^7^, additional data comprising small caliber fibers not heretofore examined, have been measured from existing electron micrographs of transverse adult optic nerve^6^. Accordingly, the fiber diameter range available for the current study is 0.41 – 2.92μm (axon diameters 0.32 – 2.57μm), compared to 0.87 – 3.13μm previously. In addition, there are three times as many fibers per mouse in the current dataset. Relative frequency histograms for axon, fiber and myelin (i.e. twice myelin radial thickness) diameters are summarized by genotype in Fig. S1, and *g* ratio histograms are summarized in Fig. S4.

The main motivation for increasing the dataset is to confirm the linearity of the previous large caliber axon-fiber diameter relation^7^ by adding small fibers close to the threshold for myelination (axons 0.18-0.20μm diameter^14^). Scatterplots of axon versus fiber diameters from three wild type (mice A-C) and three *rsh* mice (D-F) are shown in Fig. 1A and 1B, respectively, and fit using Deming regression (fits using nonlinear regression are virtually indistinguishable). For both genotypes, the Pearson’s correlation coefficients exceed *r_xy_* = 0.99, which are unchanged compared to the previous study^7^. Moreover, Box-Cox plus biological properties tests (Figs S2 and S3) strongly support linear relations for both genotypes.

**Figure 1.**
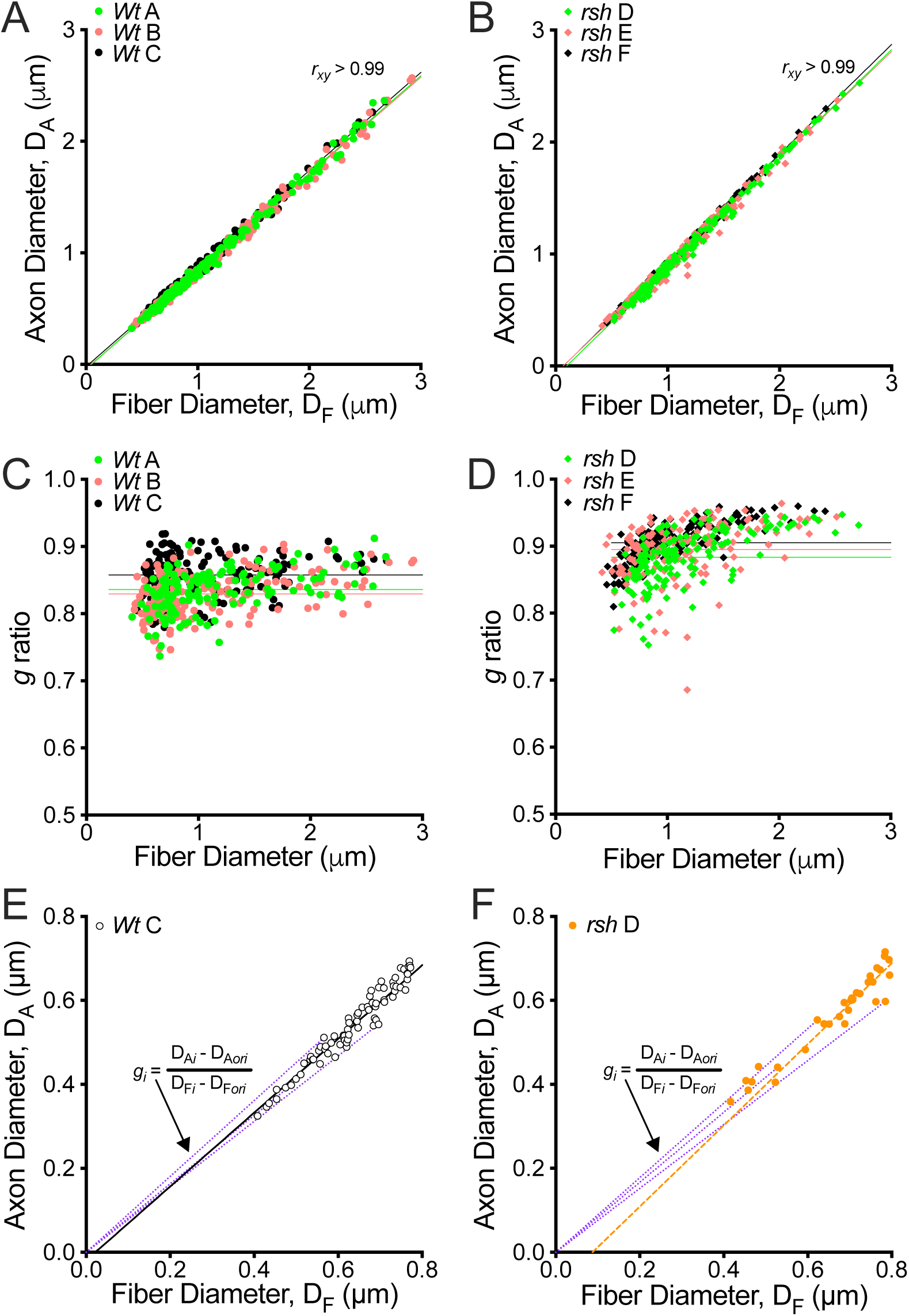
the g ratio is a slope estimator of the axon-fiber diameter plot. **A,B.** scatterplots showing axon diameter, D_A_, as a relation of fiber diameter, D_F_, measured from electron micrographs of transverse optic nerve for three wild type (*Wt* A-C) and *rsh* (D-F) mice, respectively. Pearson’s coefficients, *r_xy_*, approximate unity, and strongly support linear axon-fiber diameter relations for both genotypes. The intercepts for the regression fits in the plots are far from the Origin for the *rsh* cohort (*x*-intercept = 0.078, 95% CI: [0.03, 0.12], P = 0.017; *y*-intercept = −0.075, 95% CI: [−0.12, −0.03], P = 0.017) compared to wild type mice (*x*-intercept = 0.037, 95% CI: [0.005, 0.070], P = 0.038; *y*-intercept = −0.033, 95% CI: [−0.061, −0.005], P = 0.038). The insets show typical *g* ratio plots as a relation of fiber diameter with horizontal regression fits^7^. **C,D.** *g* ratio plots for the data in panels (A) and (B), respectively, with horizontal regression fits. These fits are consistent with a recent analysis of *rsh* optic nerve *g* ratio data^7^. The distribution of the wild type *g* ratios on the vertical axis is relatively symmetric about the regression fits (C). On the other hand, the corresponding *rsh* data are markedly skewed (D). Relative frequency histograms are shown in Fig. S4. **E,F.** Zoomed views of the plots in panels (A) and (B), respectively, show individual datapoints for single mice (panel (E), wild type mouse C; panel (F), *rsh* mouse D) in the vicinity of the Origin and highlight their connection to the regression fit (solid line). The purple dotted line segments connect several datapoints to the Origin (*ori*), the slopes of which represent *g* ratios, as demonstrated by the equation. In (E), the regression fit passes very close to the Origin so the *g* ratios approximate its slope within experimental measurement errors. In (F), the *g* ratios are poor estimates because they all approach the regression fit from the left and clearly underestimate its slope.

A second motivation is to combine large and small caliber fibers to determine if the axon-fiber diameter relation is consistent with direct proportionality, at least for the three wild type mice (Fig. 1A). Indeed, the *y*-intercepts pass very close to the Origin indicating that this important relation (i.e. the axomyelin unit model) likely holds. For *rsh* mice, the axon-fiber diameter relation was previously shown to be linear but not directly proportional (i.e. regression fits were offset from the Origin). The current result is consistent with this finding (Fig. 1B). Third is to determine if the *g* ratio plots (Fig. 1C and 1D, respectively) are similar to those in the previous study^7^ and can be fit using horizontal regression. Indeed, the larger datasets do not substantially alter the plots average *g* ratios. Thus, the data collection methods and processing protocols are consistent between studies.

In accordance with the linear axon-fiber diameter regression fits in Fig. 1A and 1B, the axomyelin unit model predicts that *g* ratio values for wild type and *rsh* mice should be independent of fiber diameter (but with different values for each genotype). And while the *g* ratios from the wild type mice are consistent with a constant *g* ratio independent of fiber diameter (Fig. 1C, also shown previously^2,5,7,15^), *g* ratios for the *rsh* cohort appear rather skewed and appear to have a somewhat positive correlation with fiber diameter (Fig. 1D), which recapitulate the results from the previous study^7^. Together, these results reveal a paradox – why do highly correlated, linear regression fits to the axon-fiber relation yield invariant *g* ratios for wild type mice, but not *rsh*?

### The g ratio plot estimates the axon-fiber diameter regression slope

This discrepancy between wild type and *rsh g* ratio plots raises questions about current understanding of the *g* ratio; both what it measures and the biological properties it represents. But an explanation emerges from consideration of the *g* ratio equation itself (eqn 4). Thus, from the standpoint of linear axon-fiber diameter plots, the *g* ratio value for each fiber is vertical rise over horizontal run, thereby defining nothing more than the slope of a line segment between two points:

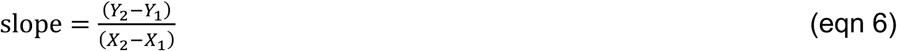

where *Y*_1_ = *X*_1_ = 0 is the Origin. This insight is highlighted in Fig. 1E, which is an expanded view of Fig. 1A near the Origin (but showing wild type mouse C only). Several line segments, *g_i_* (purple dotted lines), connect the datapoints, 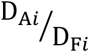, with the Origin, 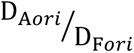. The linear regression fit (solid line) passes through the middle of the axon-fiber diameter datapoints and extrapolation passes very close to the Origin. Thus, each *g* ratio is approximately aligned with the regression fit. And because some *g* ratios are steep while others are more shallow (for constant fiber diameter, axon diameter and myelin thickness vary along the internode^16–19^), the grand average *g* ratio is a simple (crude) estimator for the regression slope ± error. Indeed, from a theoretical perspective (i.e. in the absence of experimental error) the grand average *g* ratio and regression slope are analogous (Supplement S2).

The purple dotted lines in Fig. 1F show *g* ratio line segments for *rsh* mouse D (expanded view of Fig. 1B). In contrast to wild type, the *rsh* regression fit crosses the *x*-axis at some distance from the Origin. Consequently, the *g* ratio line segments connect the Origin to datapoints from the left hand side and all slopes are more shallow than the regression fit. Thus, *g* ratios poorly estimate the axon-fiber diameter relation because they systematically underestimate the regression slope.

The purple dotted lines for *rsh* mouse D (expanded view of Fig. 1B) in Fig. 1F contrast with wild type because the regression fit crosses the *x*-axis at some distance from the Origin. Consequently, the *g* ratio line segments (dotted purple lines) connect the Origin to datapoints from the left hand side and all slopes are more shallow than the regression fit. Consequently, *g* ratios poorly estimate the axon-fiber diameter relation because they systematically underestimate the regression slope.

### Underestimation errors reflect g ratio skewness, not myelin biology

From a broader perspective, underestimating the regression slope causes data skewing, which is readily apparent in relative frequency histograms (Fig. S4). Thus, the wild type *g* ratio plots (Fig. 1C) are roughly symmetric on the vertical axis (the horizontal axis in Fig. S4A), and can be fit using a Gaussian equation. In contrast, the asymmetric *rsh* data are left skewed (Fig. 1D, long tail points down), and can be fit using a Gumbel function (eqn 1) in Fig. S4B. Moreover, averaging each of the Fig. S4 histograms and using Akaike’s Information Criterion (AICc) to compare Gaussian versus Gumbel fits to each cohort, shows there is a 99.99% probability of correctness that the wild type distribution is Gaussian, and a 99.96% probability of correctness that the *rsh* cohort is left skewed (long tail to the left).

The implications of underestimation errors are more directly revealed in Fig. 2. The *g* ratio plot in Fig. 2A shows raw data from wild type mice A-C on a large scale (gray “+” values, from Fig. 1C). The overlaid black circles are the average (mean ±95% CI) predicted *g* ratios from the individual regression fits. They have been fit with a reciprocal function (eqn 5), which is the correct equation for non-zero slope or curvilinear data (Supplement S4). The regression slope of the axon-fiber diameter plot from Fig. 1A is also shown (horizontal dotted magenta line). This line overlaps the 95% CIs for the reciprocal function, suggesting there is no statistical difference between the curvilinear fit and the regression slope from Fig. 1A, and therefore minimal underestimation errors. Nevertheless, the conspicuous curvilinearity increases asymptotically to the right and approaches (i.e. within 1%) the axon-fiber diameter regression slope for fibers between 5-10μm diameter.

**Figure 2.**
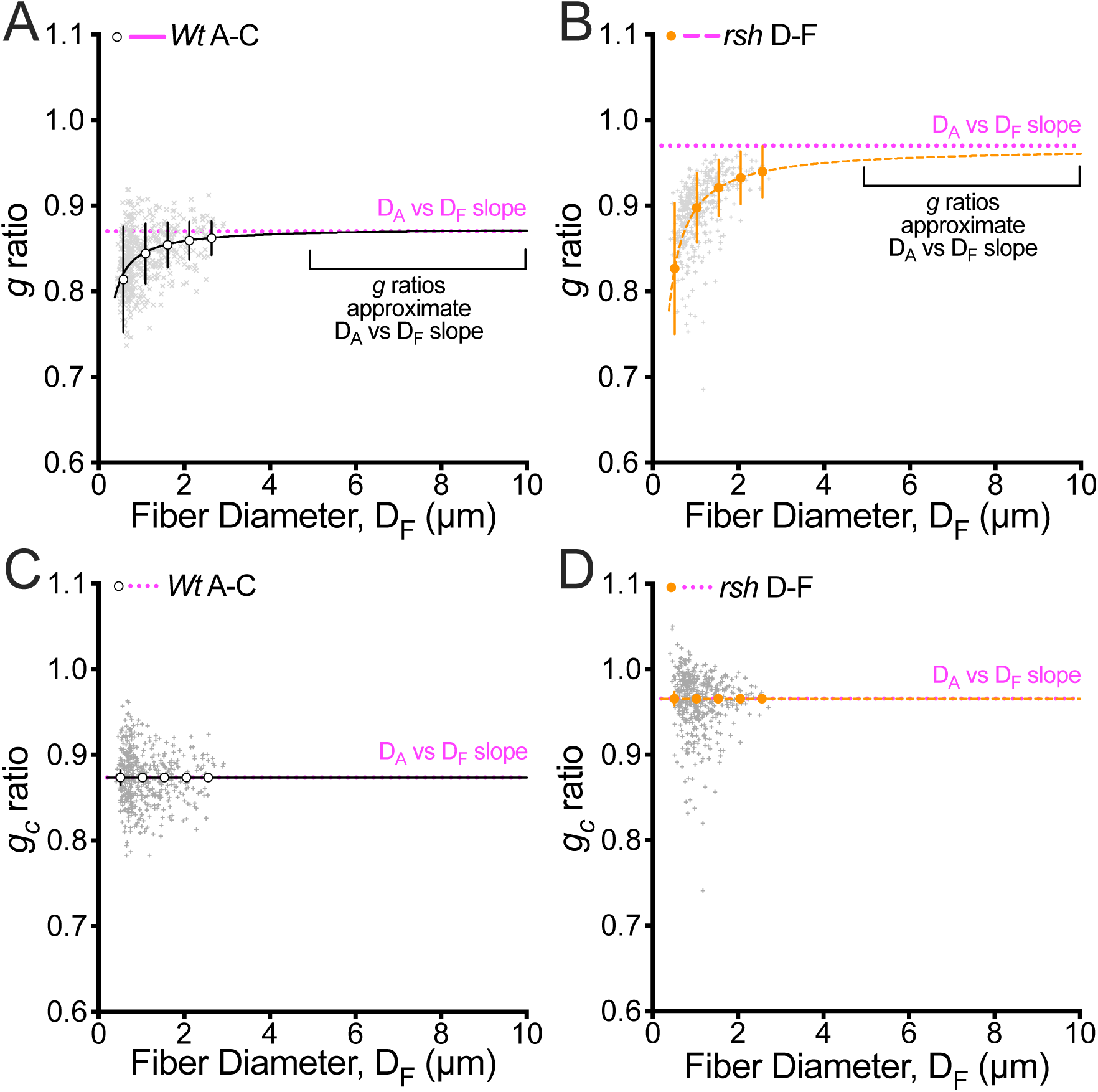
non-zero slope and curvilinear fits on g ratio plots are artifacts, not biology. **A,B.** Raw *g* ratios from wild type (A) and *rsh* (B) mouse data in Fig. 1C and 1D, respectively, are aggregated (gray “+”) to show the overall data structures across a wide range of fiber diameters. For each mouse, a reciprocal function has been fit to the data and predicted *g* ratio values averaged to generate the black and orange datapoints. These datapoints have been fit with a reciprocal function (black and orange dashed fits) to show cohort averages (means ± 95% CIs). The slopes of the axon-fiber diameter regression fits (dotted magenta lines) have been added to demonstrate the extents to which *g* ratios underestimate the actual D_A_ vs D_F_ slope. While the *g* ratio 95% CIs overlap the D_A_ vs D_F_ slope in (A), there is significant divergence for *rsh g* ratios in (B). The extrapolated reciprocal fits from the wild type and *rsh* data converge with the D_A_ vs D_F_ slope between 5-10μm fiber diameters. Historically, a number of early studies noted that *g* ratios became independent of fiber diameters above 5-10μm, but this finding is controversial^41^ (Supplement S7). **C,D.** The data in Fig. (1A) and (1B), respectively, have been used to compute *g_c_* ratios (eqn S15) in contrast to eqn S7, which at least partially account for the effects of non-zero *y*-intercepts on regression slope. The raw data are roughly symmetric on the vertical axis and accord with the slopes from the axon-fiber diameter plots.

A reciprocal function fit to the *rsh* data in Fig. 2B (gray “+” values, from Fig. 1D) shows the curvilinearity is more extreme. The magenta line is beyond the 95% CIs, indicating a statistically significant underestimation of the axon-fiber diameter regression slope. The discrepancy is pronounced for small fibers, while for large fibers the *g* ratios better approximate the axon-fiber diameter regression slope. Again, the reciprocal fit approaches the magenta line asymptotically, and is within 1% of the axon-fiber diameter regression slope for fibers at approximately 10μm in diameter. In this case, it is clear that the asymptote is the more accurate estimate. Thus, computing the grand average *g* ratio from such skewed data, as is performed in many published studies, is a substantial underestimate of the axon-fiber diameter slope.

An alternative method to *g* ratios for assessing the axon-fiber diameter relation is to compute corrected *g* ratios, (*g_c_* ratios), which at least partially account for the effects of non-zero *y*-intercepts on regression slope (Supplement S5 and S6) as shown in Fig. 2C and 2D. In contrast to the asymmetry across the magenta line in Fig. 2A and 2B, the raw datapoints (gray “+”) computed as *g_c_* ratios are symmetric. Moreover, the predicted *g_c_* ratio data and regression fits (black and orange) are coincident with the axon-fiber diameter regression slopes. These plots demonstrate directly that curvilinear *g* ratio regression fits are caused by non-zero *y*-intercepts. Thus, *g_c_* ratio plots have horizontal regression fits that are concordant with the linear axon-fiber diameter relation and the axomyelin unit model. But an outwardly disquieting and non-intuitive feature of *rsh g_c_* ratio plots emerges in Fig. 2D, where some datapoints for small caliber fibers exceed unity. This is not a miscalculation but rather, a property of the *g_c_* ratio equation, and may be associated with experimental error that becomes apparent because the *g_c_* ratios are close to unity.

### A more direct measure of the axon-fiber diameter relation – the g’ cline

Given a choice of metrics to determine the axon-fiber diameter relation, the *g* ratio is a fair estimator if, and only if, the relation is directly proportional and experimental measurement error is absent. However, the linear regression slope of the axon-fiber diameter relation is an excellent estimator under physiological and a variety of non-physiological conditions (Fig. 1), because it accounts for correlation and covariance between the axon and fiber diameters and eliminates the impact of non-zero *y*-intercepts (Supplement S2). In this regard, the regression slope (also analogous to the first derivative of eqn 3) of the axon-fiber diameter relation is parsimonious with the axomyelin unit model. This slope is herein designated the *g’* cline (pronounced gee-kl-eye-n).

To characterize properties of the *g’* cline in greater depth, datapoints from wild type mouse C and *rsh* mouse D (from Fig. 1A and B) are plotted together in Fig. 3A, with the averaged regression lines for each cohort and the equality line for unmyelinated axons (blue dotted line, D_A_ = D_F_, slope = 1). The CNS in *rsh* mice is hypomyelinated during development as a result of metabolic dysfunction^20–22^; thus, the regression fit (and *g’* cline in the equation) is intermediate between the wild type regression fit and the equality line. Further, because the *rsh* regression fit intersects the *x*-axis at some distance from the Origin, it intersects the wild type fit at an equivalence point (D_F_ = 0.44, inset in Fig. 3A). This point delineates the caliber of *rsh* fibers that are morphologically, if not functionally, comparable to their wild type counterparts. Above this caliber, *rsh* fibers are hypomyelinated compared to the controls, which accounts for more than 90% of retinal ganglion cell axons^23^.

**Figure 3.**
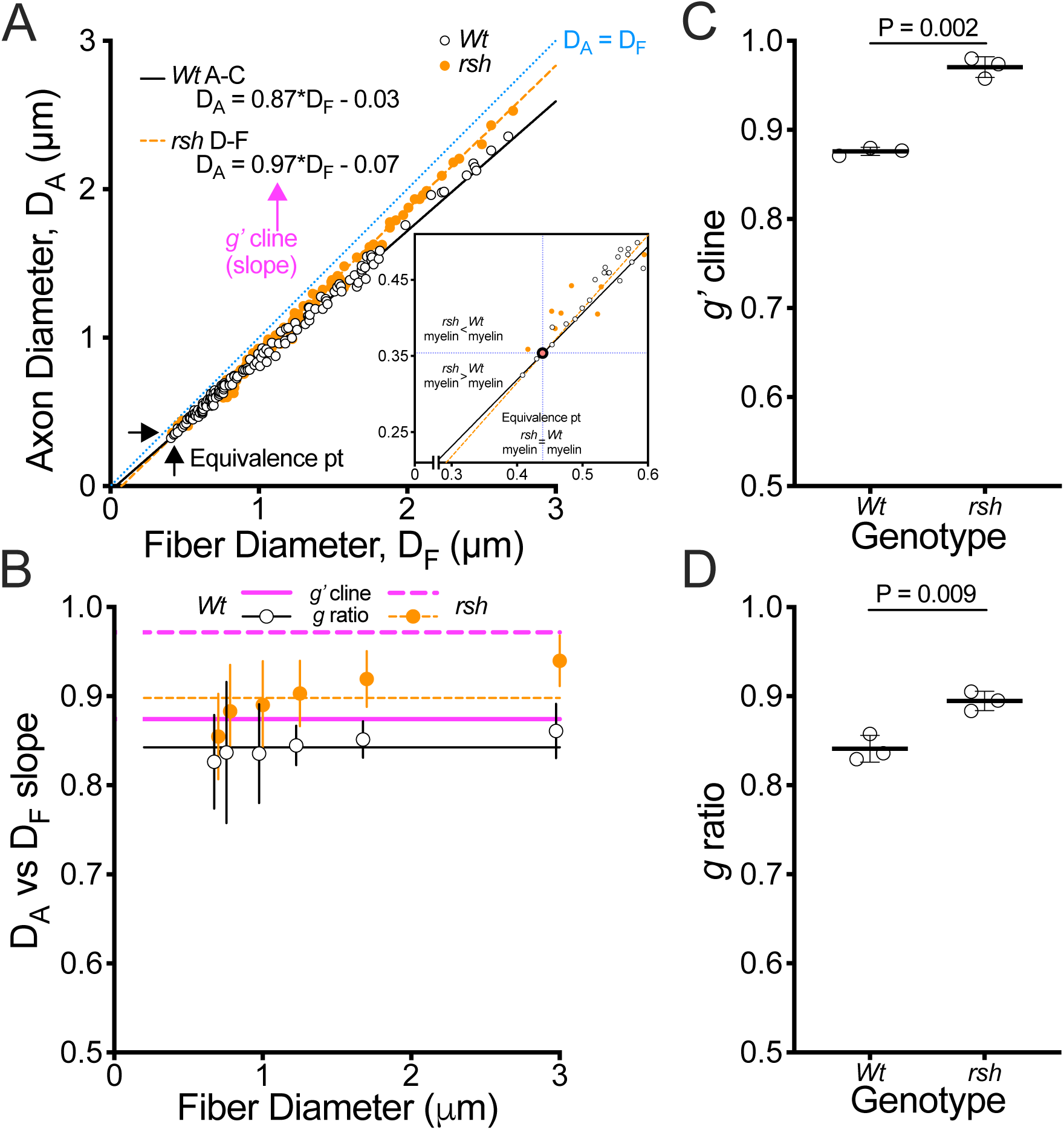
the g’ cline is a superior estimator of the axon-fiber diameter relation. **A.** combining scatterplots of the axon-fiber diameter relation for wild type (*Wt*) mouse C and *rsh* mouse D for direct comparison. The equality line (blue dotted) represents unmyelinated axons, with slope = unity defining the maximum regression slope under normal physiological conditions. The equations quantify the slopes and *y*-intercepts for the respective regression fits, including the proposed *g’* cline estimator. Because the intercepts for the *rsh* mouse are far from the Origin, and its slope is steeper than for the wild type mouse, the regression fits intersect at an equivalence point. The inset zooms in on this region to emphasize its potential significance for myelin thickness in internodes far from the equivalence point. **B.** Data from Fig. 1A and 1B processed through the analysis pipeline^7^ to define *g* ratios for the wild type and *rsh* mouse cohorts (means ±95% CI). Wild type datapoints are offset to visualize the *rsh* CIs. This plot is comparable to that reported previously for the smaller dataset, including the apparent correlation between *g* ratio and fiber diameter for the *rsh* cohort (i.e. non-zero slope of an imaginary line drawn through the datapoints). But, because the axon-fiber diameter relation is linear for the wild type and *rsh* cohorts (Figs 1A, 1B, S2 and S3), the *g’* clines are constant and have zero slope (i.e. *g’* cline is independent of fiber diameter). Thus, the *g’* cline demonstrates that the apparent non-zero slope for the *rsh g* ratio plot is an artifact. In addition, this plot emphasizes the impact of non-zero *y*-intercepts for the *rsh* cohort (shown in Fig. 2B), where the *g* ratio and *g’* cline are widely divergent. **C,D.** Welch’s *t-*tests comparing wild type and *rsh* cohorts using *g’* clines (C) or *g* ratios (D). Both metrics are highly statistically significant, but *g’* clines have greater sensitivity to detect such pathological changes.

The plot in Fig. 3B summarizes the cohort datasets from Fig. 1A and 1B after processing through the *g* ratio pipeline^7^ (summarized in Fig. S5). The *g’* clines in the equations in Fig. 3A are overlaid on this plot (magenta lines). The average *g* ratio for the wild type cohort (solid lines) reasonably approximates the *g’* cline (i.e. within 4%), which lies within most of the 95% CIs. However, these measures differ substantially for the *rsh* cohort (average *g* ratio is 92% of the *g’* cline) because of the systematic underestimation of the *g’* cline. Indeed, only the 95% CI for the 3μm diameter fibers overlap the *g’* cline. Such poor correspondence likely arises because *g* ratios conflate the effects of slope and *y*-intercept, while *g’* clines only reflect the slope. Welch’s *t*-tests for the *g’* clines (Fig. 3C) and *g* ratios (Fig. 3D) show large statistical differences between the wild type and *rsh* cohorts, although *g’* clines are more sensitive than *g* ratios.

### Interpreting axon-fiber diameter plots

A focus on axon-fiber diameter plots affords opportunities for fresh perspectives in presenting and interpreting morphological changes in internodal myelin. For example, Fig. 4A reproduces the averaged regression fit for the wild type cohort (from Fig. 3A, black line), while the *rsh* cohort is represented in Fig. 4B (dashed orange line). Because fiber diameters are the sum of axon and myelin measurements, axon diameters contribute to both the *x*- and *y*-axes. This is emphasized by parsing fiber diameter as the sum of its components in Fig. 4A and 4B. Consequently, the equality line in these plots is the D_A_ component, and D_M_ is represented by rightward horizontal shifts (i.e. red arrows) determined from the linear equations for D_M_ (Fig. 4C). The equality line-red arrow combination is the axon-fiber diameter regression fit in each plot.

**Figure 4.**
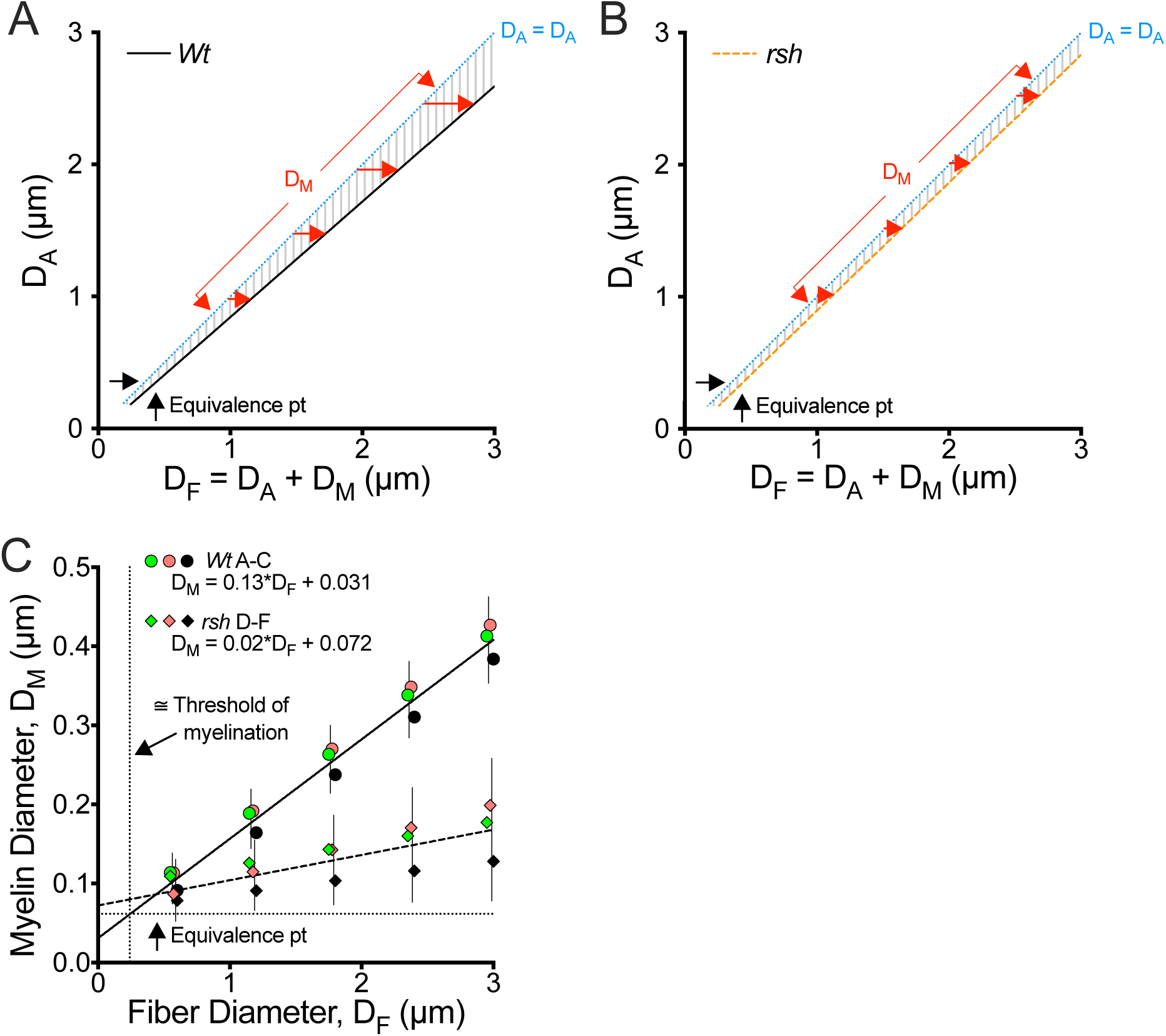
interpreting axon-fiber diameter plots with new perspective. **A,B.** axon-fiber diameter regression fits and the equivalence point from Fig. 3A, separated by genotype; wild type (*Wt*, black line) and *rsh* (dashed orange line). The *y*-axis is labeled according to axon diameter, D_A_, and the *x*-axis as D_F_, which includes the sum of its components, D_A_ and myelin diameter, D_M_ (twice the myelin radial thickness). Four examples of D_M_ are shown as horizontal red arrows. From this perspective, the equality line (blue dotted) is the D_A_ component on the *x*- and *y*-axes, which serves as a frame of reference for the axon-myelin relation. In (A), this well-known relation is apparent from the increasing width of the hatched area between the equality line and the regression fit (D_M_ is ∼14% of D_F_). In contrast, (B) this relation is lost in *rsh* oligodendrocytes, likely because of metabolic pathology, so the width of the hatched area is essentially invariant with increasing fiber diameter. **C.** values of myelin thickness (colored symbols) predicted from linear regression fits of the myelin-fiber diameter data for each mouse. The solid and dashed lines are the average regression fits for each cohort ±95% CI, which intersect at the equivalence point, consistent with Fig. 3A. As expected, the regression fit of the wild type cohort passes through the myelination threshold point (dotted vertical and horizontal lines), while the *rsh* regression fit suggests myelin is on average thicker than wild type for the smallest axons.

In Fig. 4A, the lengths of red arrows in the hatched region are proportional to fiber diameter (comprising ∼14% of the fiber diameter for all calibers), which is consistent with the known internodal property: thicker myelin around larger axons. This perspective provides a direct interpretation of the *g’* cline as the contribution of myelin diameter to the total caliber. In contrast, the *rsh g’* cline in Fig. 4B has a similar slope to the equality line, indicating that myelin thickness is constant and essentially independent of fiber caliber (i.e. red arrows are of approximately equal length). This is confirmed by the myelin-fiber diameter plot in Fig. 4C (almost a horizontal regression fit), and is consistent with previous findings in *rsh* mice CNS^8,24^.

In small fibers, the myelin diameter contributes around 10% to the total caliber but drops to only 5% for large fibers, so they are the more severely hypomyelinated. In addition, the independence of myelin thickness on fiber caliber accounts both for the non-zero *y*-intercept in Fig. 1B and the equivalence point, below which *rsh* myelin may be thicker than normal. Together, the well-characterized *rsh* myelination defect can be defined in terms of an increase in the *g’* cline and a concurrent loss of direct proportionality in the axon-fiber relation.

Hatching between the equality curve and the axon-fiber diameter regression fits in Fig. 4A and 4B demarcate regions that are proportional to the myelin-fiber diameter relation. Thus, a ratio of areas between the curves for *rsh* and wild type, respectively, should provide an estimate of the relative myelin thickness in *rsh* optic nerve, which approximates 42% of normal. The areas under the curves in Fig. 4C are analogous to the hatched regions and estimate myelin in *rsh* as 31% of normal. These estimates are roughly comparable to each other and to published estimates based on in situ hybridization and electron microscopy at 40-60% of normal^8,12,24–26^.

Finally, Fig. 4C includes vertical and horizontal dotted lines which delineate the approximate threshold diameter, below which myelinated axons are not observed^27^. In the vicinity of this quasi-boundary, the cross-sectional dimensions of wild type internodes are D_F_ ≅ 0.24μm with two wraps of myelin (0.03μm per wrap), which accords with the regression fit in Fig. 4C; thus, D_A_ ≅ 0.18μm. For *rsh* mice, fibers near this boundary may have three myelin wraps (a 50% increase in myelin thickness), with D_A_ ≅ 0.17μm. Thus together, these plots can be parsed to reveal detailed multi-faceted aspects of biology, and are relatively easily interpreted.

## Discussion

Until the 1990s, *PLP1* gene mutations in Pelizaeus-Merzbacher patients and animal models thereof were considered to cause developmental disease in the sense that immature oligodendrocytes failed to fully differentiate and achieve a myelinating state^28,29^. Subsequent studies demonstrated that the disease, while developmental, arises from poor or arrested myelinogenesis triggered by metabolic stress and resulting in apoptosis of mature oligodendrocytes^21,30^. Several studies have developed hypotheses that non-cell autonomous signals in the CNS aimed at promoting myelination (e.g. growth or differentiation factors) may actually aggravate the metabolic stress and exacerbate disease severity^31,32^.

In this regard, the current axon-fiber diameter analysis of *rsh* mice in Fig. 3A provides supporting evidence, by revealing heretofore unknown structural changes to internodal myelin in small fibers that may motivate more detailed studies. Thus, while most fibers in *rsh* optic nerve are hypomyelinated, those fibers are larger than the equivalence point and myelin thickness is constant (i.e. independent of fiber diameter^8^). This suggests most oligodendrocytes operate at maximum myelinating capacity because of the negative impact of metabolic stress^33^. In contrast, a small population of *rsh* oligodendrocytes below the equivalence point may have capacity to achieve supranormal myelin wrapping (as much as one additional wrap, Fig. 4C).

Such potential hypermyelination hints at two plausible mechanisms. On the one hand, disease-counteracting epigenetic signals drive oligodendrocytes to maximize rates of myelin synthesis. And while oligodendrocytes ensheathing small axons can respond by increasing myelin wraps, metabolic stress is triggered in those myelinating large axons, which curtails function and reduces myelin synthesis. On the other hand, the paucity of the PLP1 protein in compact myelin^21,26^ may disrupt passive membrane properties of the myelin triggering increased synthesis around small axons. But as above, metabolic stress counterposes additional myelin synthesis around larger axons.

In addition to novel insights on the impact of metabolic stress on *rsh* oligodendrocyte metabolism and myelinogenesis, the current study provides the first mathematical critique of the *g* ratio plot. In short, the *g* ratio is an inferior estimator for the *g’* cline. While they can yield satisfactory results for internodes that comport with the axomyelin unit model (axon-fiber diameter plot with zero *y*-intercept), non-zero slope or curvilinear *g* ratio plots must be fit using a reciprocal function to be meaningfully interpreted (i.e. the asymptote has defined meaning). Indeed, it is the asymptote of this function, not the mean or median *g* ratio, that best estimates the *g’* cline (Supplement S4).

Indubitably, the smallest fibers contribute most to *g* ratio curvilinearity, which is of little surprise because measurement artifacts are more significant for fibers with only two or three myelin wraps. In this regard, lack-of-resolution is the likely cause of the curvilinear fit in Fig. 2A. Although within experimental error (i.e. the 95% CIs), the artifact is apparent because electron micrographs used for the current study are 9nm / pixel. This resolution is marginally below the typical 10 – 12nm myelin period in aldehyde-fixed tissue^34^ but, optimally, should be < 3nm / pixel^12^.

An alternative to the *g* ratio plot is the *g_c_* ratio plot, which at least partially compensates for the effects of non-zero *y*-intercepts on the regression slope (Fig. 2). However, the range of *g_c_* ratio values can be unstable because they are not confined between zero and one. This counterintuitive property violates a major perception about *g* ratios, that the diameter of an axon cannot exceed the diameter of its fiber (0 < *g* ratio < 1). But, this perception arises only because our understanding of the *g* ratio is incomplete. A *g* ratio is not an assertion about individual axon and fiber diameter ratios. Rather, a set of *g* ratios, collectively, estimate the *g’* cline (and ultimately the constant of proportionality), which represents the rate of change of axon diameter with fiber diameter (analogous to a velocity-time plot in physics).

These revelations notwithstanding, the current study also demonstrates that the *g’* cline is a more general and superior measure of the axon-fiber diameter relation than has, hitherto, been proposed. Indeed, the major advantage of the *g’* cline is in its separation of the rate of change of myelin thickness from other factors, which simplifies interpretation. By focusing the quantitative analysis on the axon versus fiber diameter plot, rather than *g* ratios, the *g’* cline avoids artifacts that have persisted in the myelin field for as long as a century (e.g. Figs S6 and S7). Further, *g’* clines are more parsimonious with the axomyelin unit model and, therefore, can be interpreted in a constrained framework with greater confidence leading to better testable hypotheses.

The *g’* cline proposed herein, is so named for three reasons: 1) to emphasize its connection to the *g* ratio (Fig. 1) which is widely used and generally has been misinterpreted for more than a century; 2) to emphasize it is the regression slope in the axon-fiber diameter plot, and; 3) to explicitly acknowledge it as analogous to the first derivative of the linear regression equation. In accordance with the axomyelin unit model and multiple studies in many adult vertebrate species (i.e. steady-state myelin sheaths), the *g’* cline demonstrates that the axon-fiber diameter relation is approximately linear under normal physiological conditions. This assertion provides a rigid framework for analysis that justifies use of the *g’* cline so that appropriate mixed-effects statistics can be used to reveal incongruities in non-steady-state conditions: normal development/aging and myelin plasticity-associated learning as well as disease states.

By opting for the more generally-applicable *g’* cline analysis, *g* ratios are arguably, obsolete. However, Figs 2 and S4 demonstrate that *g* ratios are useful for exploratory data analysis^10^ by visually detecting measurement artifacts and skewness. In particular, the regression curve in Fig. 2A demonstrates the sensitivity of *g* ratios to skewness where, statistically, there is minimal likelihood of skewing and yet curvilinearity is clearly apparent. Of course, many equations have been developed to assess skewness – from Pearson’s third central moment to the commonly used *γ*_$_ (gamma) statistic – but these methods generate simple numbers, do not reliably quantify skewness, and cannot convey the visual clarity provided by graphs. Thus, *g* ratio plots remain an important tool for characterizing internodal myelin.

As an aside, when Bear & Schmitt^35^ developed the mathematics to quantify and interpret myelin birefringence, they designated the axon:fiber diameter ratio as the variable, “g”, in their equations (reviewed in^13^). This appears to be an arbitrary assignment – letters assigned alphabetically as labels for variables/constants when developing systems of equations – which has been customary practice since the Ancient Greeks. However, F.O. Schmitt was proficient at developing mathematical frameworks and adopting physics technologies to address quantitative questions in biology^36,37^. It is tempting to speculate he may have chosen “g” to symbolize the axon-fiber diameter ratio as a simple estimator for Pearson’s *γ*_1_, thereby acknowledging *g* ratio plots to assess skewness.

In two recent studies^4,7^, the axomyelin unit model was defined, an analytical pipeline proposed and experimental and statistical artifacts described, to promote more rigorous analysis and interpretation of *g* ratios in CNS and PNS fiber tracts under physiological conditions. The current article suggests an alternative approach may be possible, using the *g’* cline. A choice between *g* ratios and *g’* clines may lead to confusion about the appropriate approach. But from another perspective, the *g* ratio pipeline is a special case of the more generally applicable *g’* cline. Insofar as the axon-fiber diameter relation is directly proportional, both methods yield similar results, and preference can be exercised (e.g. depending on the need for consistency with previous studies).

From an historical perspective over the last 120 years, interpretation of *g* ratios has evolved considerably. Initially, the ratio concept was used to express the constancy of the strong linear relationship between axons and myelin sheaths in the PNS from key vertebrate species^15^. But over subsequent decades, interpreting the *g* ratio has become nuanced and complex, largely associated with the introduction of various experimental and statistical artifacts^2,3^, as well as a lapse in appreciating mathematical assumptions that justify the *g* ratio equation. Indeed, Fig. S6 shows how early these lapses arose, with examples in three landmark studies ^38–40^. Rolling back such nuances has been recently proposed for the *g* ratio plot^4,7^ within the context of the axomyelin unit model, which is derived from the *g* ratio equation itself but provides a framework to constrain the interpretation of internodal myelin properties.

A legacy observation favoring nonlinear *g* ratio plots is that *g* ratios are independent of fiber caliber for large caliber fibers (≥ 6μm, depending on the study), but positively correlated below this range^38–41^. Insight from Fig. 1E, 1F, 2A and 2B leads to a satisfying argument against this view because small fiber *g* ratios can substantially underestimate the *g’* cline, which is exacerbated by larger non-zero *y*-intercepts in the axon-fiber diameter relation (inversely proportional relation; Supplement S7, Table S1, Fig. S8). Thus, at least in part, it is a conflation of regression slope estimates and non-zero *y*-intercepts that accounts for curvilinearity in *g* ratio plots. Further, the notion that *g* ratios are constant only for large diameter fibers is dispelled (Supplement S7).

A corollary to the view that *g* ratios and fiber diameters are correlated is that the axon-fiber diameter relation is nonlinear. However this too is unlikely (Fig. 4). Thus, when fiber diameter is parsed into its two components, D_A_ + D_M_, the myelin component is small in comparison to the axon, and its rate of increase is less than 20% of that for the axon (Fig. 2A, *Wt* myelin slope = 0.13, Fig. 4C; *Wt* axon slope = 0.87). Thus, the strong axon-fiber diameter correlation is largely driven by the axon diameter equality line, which is by definition linear and is also consistent with the Box-Cox analysis in Figs S2 and S3. Nevertheless, the possibility of mild nonlinearity in the myelin-fiber diameter relation, as suggested by Hildebrand and colleagues^34,42^, cannot be completely excluded and underscore some of the problems of linear regression used as a statistical test to compare *g* ratio or axon-fiber diameter slopes. More sophisticated analyses such as mixed-effects modeling are necessary to dissociate such correlated measurement artifacts from biological changes.

Finally, in view of the vulnerability of *g* ratio measurements to experimental artifacts, their use in quantifying internodal myelin properties in PNS and CNS seems precarious. Afterall, pioneers like Francis Galton, Karl Pearson and others developed the statistical mathematics for linear regression prior to the twentieth century^43^, decades before the *g* ratio was popularized. Such advances notwithstanding, crucial technology was missing at that time – the computer – and processing hundreds or thousands of datapoints with complex equations was exceeding tedious, and error-prone. With only log tables and graph paper, simple graphics-based shortcuts were often used to analyze data^44^. But since the 1990s, statistics software has become ubiquitous. Perhaps the era of the *g* ratio as a quantitative tool has passed, and like other error-prone ratio-based methods such as Scatchard and Lineweaver-Burk plots, should go the way of the Dodo.

## Acknowledgements

Thank you to Cherie Southwood for her technical expertise and dedication to excellence, and discussions with colleagues: Drs Audrey Wu, CMMG, Wayne State University; Anne Boullerne, Department of Anesthesiology, University of Illinois at Chicago, Chicago, IL; Douglas Feinstein, Department of Anesthesiology, University of Illinois at Chicago, Chicago, IL; and Jeff Dupree, Department Anatomy & Neurobiology, Virginia Commonwealth University, Richmond, VA. This work was supported by the Gow/Southwood Trust and to the National Multiple Sclerosis Society (RG-2305-41469).

