## Supplementary material for "Demystifying The Myelin *g* ratio: Its Origin, Derivation and Interpretation": Gow_A_Demystifying_the_myelin_g_ratio_Suppl.pdf

DATA SUPPLEMENT

Running Title: Demystifying the *g* ratio

Alexander Gow[1,2,3,*]

[1] Center for Molecular Medicine and Genetics, [2] Dept of Pediatrics, [3] Dept of Neurology, Wayne State University School of Medicine, Detroit, MI, 48201, USA.

*** Correspondence:** Dr Alexander Gow
Center for Molecular Medicine and Genetics,
3216 Scott Hall, 540 E Canfield Ave,
Wayne State University School of Medicine,
Detroit, MI, 48201.

**#Figs, #Suppl Figs, #Tables, #Suppl Tables, #Pages:**

**Key words:** myelin, axon, myelinated fiber, *g* ratio, nervous system, white matter

Supplemental Material

The seven supplemental sections below comprise algebraic derivations defining a number of properties and drawing parallels between different aspects of the axomyelin unit model, as well as providing the underlying principles of artifacts that are commonly misinterpreted as biological properties of the myelin internode. These sections are more-or-less standalone and are referred to in different sections of the main article, as necessary. They provide further explanation for the interested reader but are not necessary for comprehending the content of the article.

*Section S1 – should the fiber diameter or the axon diameter be expressed on the x-axis?*

Initial exploration of the structural and functional characteristics of axons and myelin sheaths typically represented fiber diameter on the *x*-axis. This practice began to change after Gasser and Grundfest[1] demonstrated that conduction velocity is correlated with axon diameter. In contemporary literature, axon diameter is almost exclusively represented on the *x*-axis. The argument in favor of fiber diameter on the *x*-axis is in part a matter of elegance in the equations describing the axon-fiber diameter relation.

The axon-fiber diameter relation is widely known to be linear and can be described by the linear equation:

$$\mathrm{D_A} = \mathrm{slope} * \mathrm{D_F} + y\mathrm{intercept} \qquad \text{(eqn S1)}$$

where, $\mathrm{D_A}$ and $\mathrm{D_F}$ are the axon and fiber diameters, respectively, $\mathrm{D_F} > \mathrm{D_A}$ and $y\mathrm{intercept} \leq 0$ under normal physiological conditions (the axomyelin unit model)[2]. To compute the *g* ratio for each axon-fiber measurement, the fiber diameter is divided into the axon diameter:

$$\frac{\mathrm{D_A}}{\mathrm{D_F}} = \frac{\mathrm{slope} * \mathrm{D_F}}{\mathrm{D_F}} + \frac{y\mathrm{intercept}}{\mathrm{D_F}} \qquad \text{(eqn S2)}$$

which can be reduced according to the value of the $y\mathrm{intercept}$:

$$\frac{\mathrm{D_A}}{\mathrm{D_F}} = \mathrm{slope} = g \text{ ratio} \qquad \text{(eqn S3a)}$$

if $y\mathrm{intercept} = 0$, or:

$$\frac{\mathrm{D_A}}{\mathrm{D_F}} = \mathrm{slope} + \frac{y\mathrm{intercept}}{\mathrm{D_F}} \quad \Rightarrow \quad \frac{\mathrm{D_A} - y\mathrm{intercept}}{\mathrm{D_F}} = \mathrm{slope} = g \text{ ratio} \qquad \text{(eqn S3b)}$$

if $y\text{intercept} \neq 0$. Equations S3a and S3b are equivalent to eqn 4 and 5, respectively, and the definition of the *g* ratio emerges naturally. Further, because $\mathrm{D_F} > \mathrm{D_A}$, the possible range of values for the *g* ratio are constrained, $0 < g\ \text{ratio} < 1$, which constitutes a well-behaved function. In contrast, if axon diameter is represented on the *x*-axis, then the linear equation is:

$$\mathrm{D_F} = \text{slope} * \mathrm{D_A} + y\text{intercept} \qquad \text{(eqn S4)}$$

where,

$$\frac{\mathrm{D_F}}{\mathrm{D_A}} = \text{slope} = \frac{1}{g\ \text{ratio}} \qquad \text{(eqn S5a)}$$

for $y\text{intercept} = 0$, and:

$$\frac{\mathrm{D_F}}{\mathrm{D_A}} = \frac{\text{slope}*\mathrm{D_A}}{\mathrm{D_A}} + \frac{y\text{intercept}}{\mathrm{D_A}} \quad \Longrightarrow \quad \frac{\mathrm{D_F}}{\mathrm{D_A}} = \text{slope} + \frac{y\text{intercept}}{\mathrm{D_A}} = \frac{1}{g\ \text{ratio}} \qquad \text{(eqn S5b)}$$

for $y\text{intercept} \neq 0$. Because $\mathrm{D_F} > \mathrm{D_A}$, the range of values for $\frac{1}{g\ \text{ratio}}$ are only partially constrained, $1 < \frac{1}{g\ \text{ratio}} < \infty$. While this incongruity is merely technical and can be easily remedied to recover the *g* ratio in the expected range, the approach is clumsy, and the equations inelegant.

*Section S2 - the g ratio is analogous to regression slope for the axon-fiber diameter relation*

From a theoretical consideration of the axomyelin unit model[2], the hypothesized direct proportionality between an individual axon diameter, $\mathrm{D}_{\mathrm{A}i}$, and the corresponding fiber diameter, $\mathrm{D}_{\mathrm{F}} = [\mathrm{D}_{\mathrm{A}}$ + (2 * myelin radial thickness, $\mathrm{D}_{\mathrm{M}}$)], can be represented as $\mathrm{D}_{\mathrm{A}} \propto \mathrm{D}_{\mathrm{F}}$. This proportionality is hypothesized as a linear function in the absence of experimental measurement error, and the constant of proportionality, $\kappa$, links $D_A$ and $D_F$ such that:

$$\mathrm{D}_{\mathrm{A}} = \kappa * \mathrm{D}_{\mathrm{F}} \qquad \text{(eqn S6)}$$

where any value for $D_F$ yields a unique $D_A$ (i.e. a deterministic system), and the point-slope is constrained, $0 < \kappa < 1$. Values for $\kappa \geq 1$ or $\kappa \leq 0$ in eqn S6 cannot represent a myelinated axon, by definition, because: $\kappa \neq 0$ is the typical assumption implied by correlated variables; $D_A$ and $D_F$ are positively correlated, and; $\mathrm{D}_{\mathrm{A}} < \mathrm{D}_{\mathrm{F}}$ (Fig. 1).

In reality, experiments involve multiple axon and fiber diameter measurements, $(\mathrm{D}_{\mathrm{F}i}, \mathrm{D}_{\mathrm{A}i})$ where $i = 1, \ldots, n$, so eqn S6 can be rearranged and expressed in terms of averaged values for $D_F$ and $D_A$:

$$\kappa = \frac{\mathrm{D}_{\mathrm{A}i}}{\mathrm{D}_{\mathrm{F}i}} = \frac{\overline{\mathrm{D}}_{\mathrm{A}}}{\overline{\mathrm{D}}_{\mathrm{F}}} = g \text{ ratio} \qquad \text{(eqn S7)}$$

where $\overline{\mathrm{D}}_{\mathrm{F}} = \frac{\sum \mathrm{D}_{\mathrm{F}i}}{n}$, $\overline{\mathrm{D}}_{\mathrm{A}} = \frac{\sum \mathrm{D}_{\mathrm{A}i}}{n} = \frac{\sum(\kappa * \mathrm{D}_{\mathrm{F}i})}{n} = (\kappa * \overline{\mathrm{D}}_{\mathrm{F}})$, and the quotient of these variables is the *g* ratio, which is an estimator for the true slope, $\kappa$.

Inextricably, experiments involve measurement error, which leads to more than one value of $D_A$ for every $D_F$, even if the two variables are directly proportional. Thus, the two variables are linked by a relation (not a function). Linear regression for this relation yields the regression slope, $m$, which is a ratio of the covariance (Cov) between $D_A$ and $D_F$, and the variance (Var) in $D_F$, expressed as:

$$m = \frac{\mathrm{Cov}(\mathrm{D}_{\mathrm{F}}, \mathrm{D}_{\mathrm{A}})}{\mathrm{Var}(\mathrm{D}_{\mathrm{F}})} = \frac{\sum(\mathrm{D}_{\mathrm{A}i} - \overline{\mathrm{D}}_{\mathrm{A}})(\mathrm{D}_{\mathrm{F}i} - \overline{\mathrm{D}}_{\mathrm{F}})}{\sum(\mathrm{D}_{\mathrm{F}i} - \overline{\mathrm{D}}_{\mathrm{F}})^2} \qquad \text{(eqn S8)}$$

where, $\mathrm{Cov}(\mathrm{D}_{\mathrm{F}}, \mathrm{D}_{\mathrm{A}}) = \frac{\sum(\mathrm{D}_{\mathrm{A}i} - \overline{\mathrm{D}}_{\mathrm{A}})(\mathrm{D}_{\mathrm{F}i} - \overline{\mathrm{D}}_{\mathrm{F}})}{n}$ is the numerator, and $\mathrm{Var}(\mathrm{D}_{\mathrm{F}}) = \frac{\sum(\mathrm{D}_{\mathrm{F}i} - \overline{\mathrm{D}}_{\mathrm{F}})^2}{n}$ is the denominator. Then, substituting eqn S7 into eqn S8 yields the analogous expression in terms of $D_F$ and $\kappa$:

$$m = \frac{\sum(\kappa * \mathrm{D}_{\mathrm{F}i} - \kappa * \overline{\mathrm{D}}_{\mathrm{F}})(\mathrm{D}_{\mathrm{F}i} - \overline{\mathrm{D}}_{\mathrm{F}})}{\sum(\mathrm{D}_{\mathrm{F}i} - \overline{\mathrm{D}}_{\mathrm{F}})^2} = \frac{\kappa * \sum(\mathrm{D}_{\mathrm{F}i} - \overline{\mathrm{D}}_{\mathrm{F}})(\mathrm{D}_{\mathrm{F}i} - \overline{\mathrm{D}}_{\mathrm{F}})}{\sum(\mathrm{D}_{\mathrm{F}i} - \overline{\mathrm{D}}_{\mathrm{F}})^2} \qquad \text{(eqn S9)}$$

which simplifies to $m = \kappa$. Thus the slope, $m$, is also an estimator for $\kappa$. With a choice between using the $g$ ratio or the regression slope to estimate the true proportionality constant for the axon-fiber diameter relation, a comparison between the two metrics reveals the latter to be a simple (i.e. inefficient) estimator because it depends only on the ratio between $D_A$ and $D_F$, and discounts the covariance and variance between them.

Under theoretical conditions, eqn S7 describes the hypothetical direct proportionality of the axon-fiber diameter relation (i.e. the axomyelin unit model). However, this relation can be modified to be of greater applicability under experimental conditions. To this end, the axon-fiber diameter relation is satisfactorily described under a variety of physiological and non-physiological conditions by the general linear regression equation:

$$\mathrm{D}_{\mathrm{A}i} = (\mathrm{slope} * \mathrm{D}_{\mathrm{F}i}) + y\mathrm{intercept} \qquad \text{(eqn S10)}$$

*Section S3 - g ratio plots under experimental conditions when $yintercept \cong 0$*

Deriving an equation to compute *g* ratios is a simple matter of dividing both sides by the fiber diameter, and eqn S7 can be recovered by assuming the *y*-intercept passes through the Origin (or is negligible, i.e. $y\text{intercept} \cong 0$). When this assumption is reasonably justifiable, the regression fit to the corresponding *g* ratio plot is linear and has zero slope (i.e. the *g* ratio is independent of fiber diameter). This can easily be understood by substituting eqn S6 into eqn S7:

$$\mathrm{D}_{\mathrm{A}i} = \kappa * \mathrm{D}_{\mathrm{F}i} \quad \Longrightarrow \quad \frac{(\kappa * \mathrm{D}_{\mathrm{F}i})}{\mathrm{D}_{\mathrm{F}i}} = g \text{ ratio} \qquad \text{(eqn S11)}$$

and assessing the function for large and small caliber fibers. Thus, the upper limit is:

$$\lim_{\mathrm{D}_{\mathrm{F}} \longrightarrow \infty} \frac{(\kappa * \mathrm{D}_{\mathrm{F}i})}{\mathrm{D}_{\mathrm{F}i}} = \kappa \qquad \text{(eqn S12a)}$$

The lower limit is actually the minimum caliber axon that is myelinated in the CNS of vertebrates, which corresponds to $\mathrm{D}_{\mathrm{F}i} \cong 0.24$[3]:

$$\frac{(\kappa * 0.24)}{0.24} = \kappa \qquad \text{(eqn S12b)}$$

Thus, eqns S7a,b demonstrate that for all values of $\mathrm{D}_{\mathrm{F}}$, the *g* ratio is completely determined by the slope, which is a constant ($0 < \kappa < 1$, eqn S6). Accordingly, a regression fit to the *g* ratio plot must be a horizontal line (i.e. with zero slope), and the *g* ratio must be independent of fiber diameter.

*Section S4 - g ratio plots under experimental conditions when $yintercept \neq 0$*

On the other hand, if the assumption, $y\text{intercept} \cong 0$, is not justifiable, then eqn S7 generates artifactual non-zero slope or curvilinear *g* ratio plots. In such cases (Figs 2A, 2B, S6 and S7), the *g* ratio plot can only be interpreted correctly if it is fit using a reciprocal function (i.e. a hyperbolic function in Euclidian space):

$$\frac{\mathrm{D}_{\mathrm{A}i}}{\mathrm{D}_{\mathrm{F}i}} = \frac{(\kappa * \mathrm{D}_{\mathrm{F}i})}{\mathrm{D}_{\mathrm{F}i}} + \frac{y\text{intercept}}{\mathrm{D}_{\mathrm{F}i}} = \kappa + \frac{y\text{intercept}}{\mathrm{D}_{\mathrm{F}i}} \qquad \text{(eqn S13)}$$

This function has several properties relevant to the myelin internode. First, the regression fit is asymptotic toward the right and second, the asymptote is the estimator for the slope of the axon-fiber diameter relation. This can be understood from the upper limit for eqn S13:

$$\lim_{\mathrm{D}_{\mathrm{F}} \longrightarrow \infty} \frac{(\kappa * \mathrm{D}_{\mathrm{F}i})}{\mathrm{D}_{\mathrm{F}i}} + \frac{y\text{intercept}}{\mathrm{D}_{\mathrm{F}i}} = \kappa \qquad \text{(eqn S14a)}$$

because the $y\text{intercept}$ is a constant, so the second term on the left side tends to zero for large caliber fibers. Only then does *g* ratio approximate the slope of the axon-fiber diameter relation (i.e. at the asymptote). Thus, eqn S14a demonstrates that computing the average or median *g* ratio from an experiment, which is routine in contemporary literature, can lead to erroneous values and false-positive or false-negative results.

An unjustified assumption that $y\text{intercept} \cong 0$ can cause additional artifacts in *g* ratio plots, which are understood from the lower limit for eqn S13, where $\mathrm{D}_{\mathrm{F}i} \cong 0.24$:

$$\frac{(\kappa * 0.24)}{0.24} + \frac{y\text{intercept}}{0.24} = \kappa + \frac{y\text{intercept}}{0.24} \qquad \text{(eqn S14b)}$$

In this case, when $y\text{intercept} < 0$ the impact on the plot leading is a rapid decrease in *g* ratios for the smallest caliber fibers, which generates an apparent non-zero slope or curvilinear correlation between *g* ratio and fiber diameter. In contrast, when $y\text{intercept} > 0$, *g* ratios for the smallest fibers are higher and the *g* ratio plot appears to have a negative correlation.

*Section S5 - non-intuitive consequences when $yintercept \neq 0$*

The *g* ratio transformation (eqn S7) is a well-behaved function (i.e. $0 < g \text{ ratio} < 1$), which has likely contributed to its preservation for describing the axon-fiber relation, and as an estimator for $\kappa$. This property is understood from eqns S7a,b where both limits are completely determined by the proportionality constant, $0 < \kappa < 1$ (eqn S6).

However, there are negative consequences for describing the myelin internode using eqn S7 under some circumstances. For example, ratioing is a nonlinear transformation, and *g* ratio datasets can become heavily skewed near the boundary at unity[4]. Further, *g* ratios are very sensitive to non-zero *y*-intercepts, leading to erroneous estimates of $\kappa$ and false positive or negative results in statistical tests.

At least for data with non-zero *y*-intercepts, some shortcomings of *g* ratios can be eliminated with a simple correction by substituting eqn S6 into eqn S12 (rather than eqn S7):

$$\mathrm{D}_{\mathrm{A}i} = \kappa * \mathrm{D}_{\mathrm{F}i} \quad \Longrightarrow \quad \text{corr.}\, g \text{ ratio} = \frac{(\kappa * \mathrm{D}_{\mathrm{F}i}) - y\text{intercept}}{\mathrm{D}_{\mathrm{F}i}} \qquad \text{(eqn S15)}$$

The corr. *g* ratio ($g_c$ ratio), does not generate skewed data distributions (compare Fig. 2A and 2B with Fig. 2C and 2D), and regression fits to $g_c$ ratio plots are horizontal in accordance with the axomyelin unit model. Nevertheless, $g_c$ ratios share some properties with *g* ratios, like the upper limit:

$$\lim_{\mathrm{D}_\mathrm{F} \to \infty} \frac{(\kappa * \mathrm{D}_{\mathrm{F}i})}{\mathrm{D}_{\mathrm{F}i}} - \frac{y\text{intercept}}{\mathrm{D}_{\mathrm{F}i}} \approx \kappa \qquad \text{(eqn S16a)}$$

while the lower limit ($\mathrm{D}_{\mathrm{F}i} \cong 0.24$) is less constrained:

$$\frac{(\kappa * 0.24)}{0.24} - \frac{y\text{intercept}}{0.24} = \kappa - \frac{y\text{intercept}}{0.24} \qquad \text{(eqn S16b)}$$

In this case, if $y\text{intercept} > 0$, then $g_c \text{ ratio} < \kappa$. However, if $y\text{intercept} \leq 0$, $\kappa - \left(\frac{y\text{intercept}}{0.24}\right) \geq \kappa$, and values for the $g_c$ ratio can exceed unity (i.e. $g_c$ ratio > 1). Such values are real (and are caused by experimental errors), albeit unexpected. But as $\mathrm{D}_\mathrm{F}$ increases, eqn S16b approaches $\kappa$ and the values for $g_c$ ratio <1. Thus, $g_c$ ratio > 1 only for the smallest diameter fibers (Fig. 2D). Technically, a sufficiently large *y*-intercept, which can arise from limit-of-resolution artifacts, may generate negative values for $g_c$ ratio.

*Section S6 - g ratio versus g' cline as an estimator for $\kappa$*

Equations S1-S4 demonstrate algebraically that the direct proportionality of the axon and fiber diameter relation (the axomyelin unit model) under theoretical conditions, is analogous to the regression slope, $m$, of the axon-fiber diameter relation under experimental conditions. Further, eqn S11 shows that the *g* ratio is also an estimator for $\kappa$, and is typically expressed either as the grand average *g* ratio $\pm$ S.D. or plotted as a relation of fiber diameter (the *g* ratio plot). However, the extreme sensitivity of *g* ratios to experimental error from non-zero *y*-intercepts in the axon-fiber diameter relation, as well as other measurement artifacts[2], makes this metric suboptimal under many experimental conditions.

Such shortcomings can be at least partially compensated using a reciprocal function fit to *g* ratio plots or by computing $g_c$ ratios. But a more direct approach is simply to use the regression slope of the axon-fiber diameter relation itself, which is designated herein as the *g'* cline (Fig. 3). In contrast to the *g* ratio, the *g'* cline is an efficient metric because it accounts for covariance between axon and fiber diameters, and variance in the fiber diameter measurements (eqn S8). Further, it is insensitive to non-zero *y*-intercepts and is more generally applicable under a variety of physiological and pathological conditions.

*Section S7 - the constancy of g ratios for large diameter fibers*

A controversial view stretching back almost a century suggests *g* ratios are constant for myelinated fibers greater than 6-10μm in diameter, but decrease progressively with caliber for smaller axons. Thus, small fibers are thought to have disproportionately thicker myelin sheaths than large fibers. This notion has been reviewed in detail[5], and argued to be unlikely based on technical errors in several influential studies.

Although the black and orange regression curves in Fig. 2A and 2B are consistent with the purported *g* ratio constancy of large fibers, this property is actually correlated with the magnitude of the *y*-intercept from the axon-fiber diameter scatterplot. This correlation can be understood by rearranging and equating eqn S10 under two conditions a and b, where $y\text{intercept}_{\mathrm{a}} = 0$ and $y\text{intercept}_{\mathrm{b}} \neq 0$, to find the intersection between the asymptote of the reciprocal function and the *g'* cline (black/orange fits with the dotted magenta lines in Fig. 2A and 2B). Thus:

$$\frac{\mathrm{D}_{\mathrm{A}i}}{\mathrm{D}_{\mathrm{F}i}} = \mathrm{m}_{\mathrm{a}} \qquad \text{(eqn S17a)}$$

$$\frac{\mathrm{D}_{\mathrm{A}i}}{\mathrm{D}_{\mathrm{F}i}} = \mathrm{m}_{\mathrm{b}} + \frac{y\text{intercept}_{\mathrm{b}}}{\mathrm{D}_{\mathrm{F}i}} \qquad \text{(eqn S17b)}$$

where $\mathrm{m}_{\mathrm{a}}$ is the $\mathrm{D}_{\mathrm{A}}$ versus $\mathrm{D}_{\mathrm{F}}$ regression slope (magenta *g'* clines in Fig. 2A and 2B) and $\mathrm{m}_{\mathrm{b}}$ is the *g* ratio corresponding with the asymptote of the reciprocal function as fiber diameter increases (black/orange curves).

Because eqn S17b is asymptotic at $\mathrm{m}_{\mathrm{b}}$, a specific value for $\mathrm{D}_{\mathrm{F}}$ can only be defined when the magenta and black/orange curves are in proximity; for example, within a tolerance, $\delta$, of 1% (i.e. $(1-\delta) * \mathrm{m}_{\mathrm{a}}$). Furthermore, at this intersection, eqn S17a,b describe the same *g* ratio; thus, $\mathrm{m}_{\mathrm{b}} = \mathrm{m}_{\mathrm{a}}$. By incorporating a $\delta$ tolerance parameter in eqn S17a and substituting for $\mathrm{m}_{\mathrm{b}}$ in eqn S17b, we arrive at an expression for the relative convergence of the curves:

$$(1-\delta) * \mathrm{m}_{\mathrm{a}} = \mathrm{m}_{\mathrm{a}} + {}^{y\text{intercept}_{\mathrm{b}}}\!/_{\mathrm{D}_{\mathrm{F}i}} \qquad \text{(eqn S18)}$$

where $\mathrm{D}_{\mathrm{F}}$ is the fiber diameter at convergence. Finally, rearranging eqn S18 yields the equation:

$$\mathrm{D}_{\mathrm{F}i} = {}^{y\text{intercept}_{\mathrm{b}}}\!/_{((1-\delta) * \mathrm{m}_{\mathrm{a}} - \mathrm{m}_{\mathrm{a}})} \qquad \text{(eqn S19)}$$

which is used to solve the wild type and *rsh* datasets (Fig. 2A and 2B) and a number of prominent studies from the literature (Table S1), and plotted in Fig. S8 to demonstrate the linear relation for $D_F$ convergence as a function of *y*-intercept. Together, this analysis provides an explanation that discounts the likelihood of constant *g* ratio only for large diameter fibers.

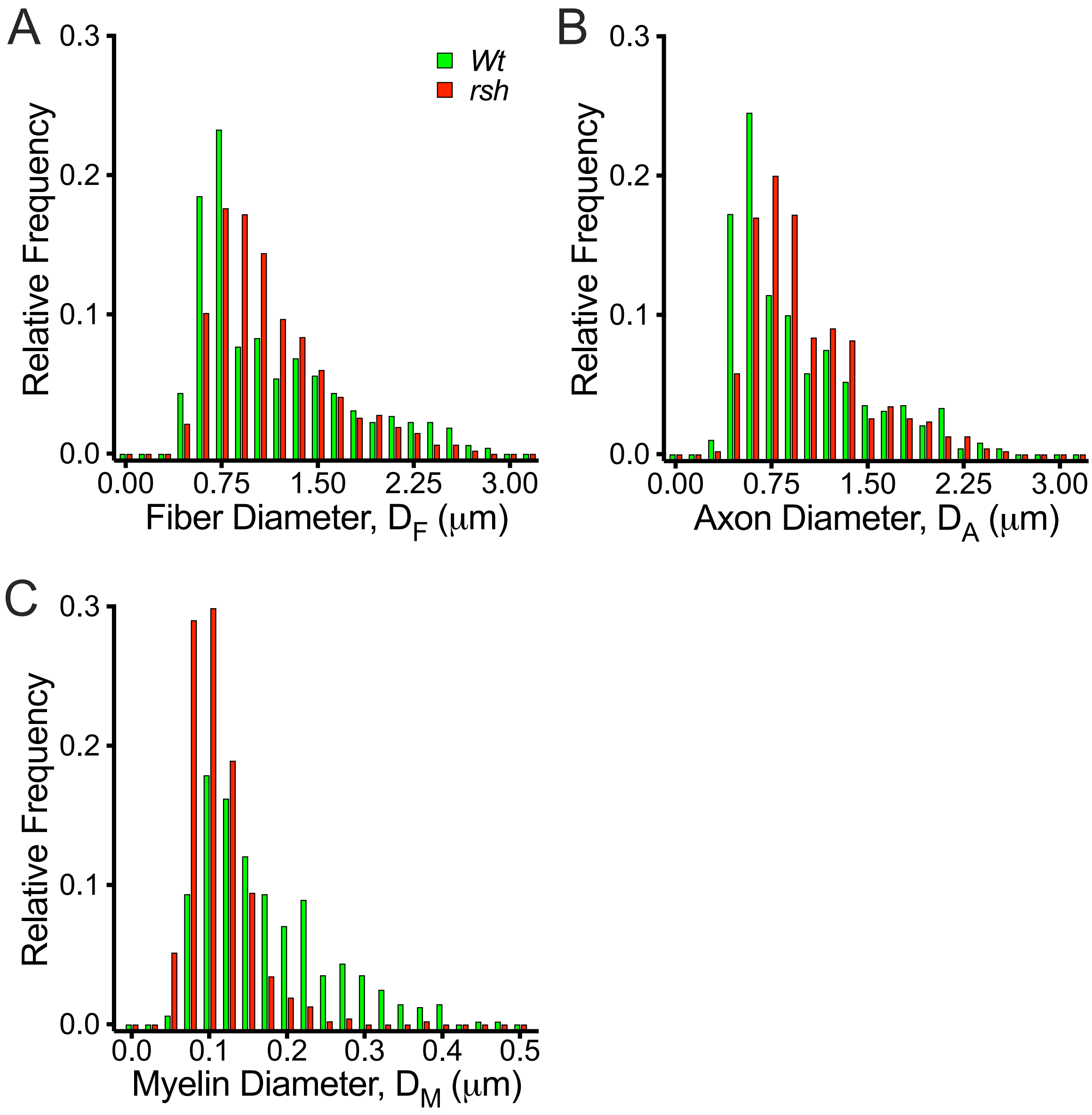


*Supplemental Figure 1 – relative frequency histograms for the current study data*

The data from each mouse in this study are combined according to genotype and interleaved (green is wild type, red is *rsh*) to generate four histograms. **A.** plot showing the distribution of fiber diameters for the wild type and *rsh* cohorts. In both cohorts, the distributions are heavily skewed to the right. The overall shape and range of fiber diameters are similar. **B.** plot showing the distribution of axon diameters. The right skewing, shape and range for both cohorts are similar and are comparable to (A) because of the strong linear relation between axon and fiber diameters. **C.** plot showing the myelin distribution, which is heavily right skewed for the wild type cohort and has a far greater range (thicker myelin around many axons) than for the *rsh* cohort. In addition, the *rsh* data are much less right skewed, suggesting myelin thickness is significantly more uniform for different caliber axons, which is consistent with Fig. S4.

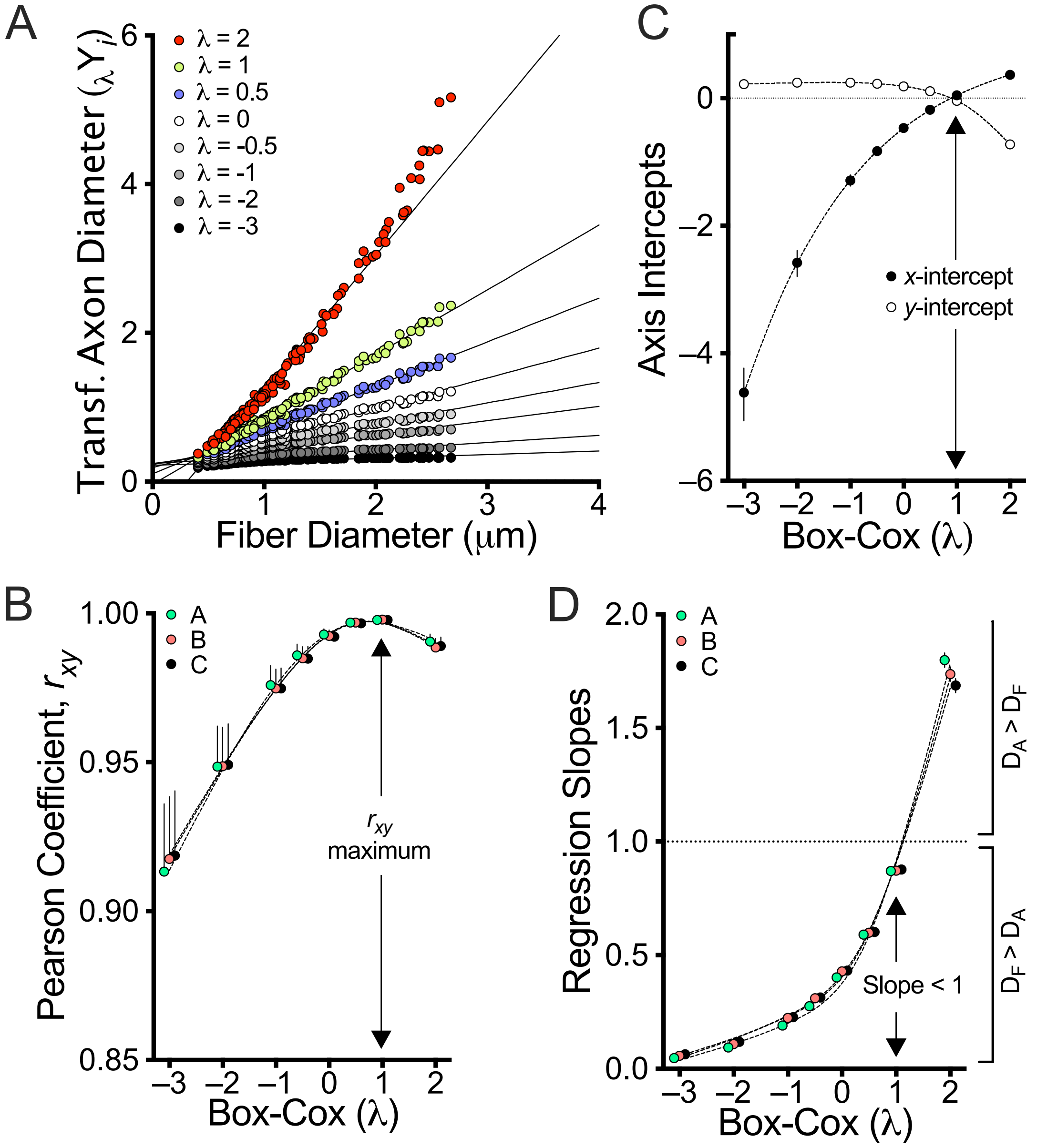


*Supplemental Figure 2 – statistical and biological constraints on wild type axon-fiber diameter plots*

**A.** Summary of a Box-Cox lambda transformation series for axon diameters from mouse C plotted against the respective fiber diameters (Fig. 1A). Each transformation is fit using linear regression to estimate parameters that identify the most likely axon-fiber diameter relation (linear versus nonlinear). **B.** Pearson's correlation coefficients for the regression fits (mice A-C) are plotted against the corresponding Box-Cox lambda values. The maximum $r_{xy}$ value, for $\lambda = 1$, is the most likely axon-fiber diameter relation as confirmed by mixed-effects analysis with Geisser-Greenhouse correction ($F_{(1.318,2.635)} = 2474$, $\varepsilon = .19$, P = <.0001) and post hoc Sidak's multiple comparison's tests show that no other values of lambda yield superior correlations (P < .007). **C.** The *y*-intercepts ±95% CI for the regression fits in (A) plotted against the Box-Cox lambda values. Biological constraints on the regression fits – *x*-intercept ≤ 0 and *y*-intercept > 0 – exclude transformations of $D_A$ where $\lambda < 1$. **D.** The constraint of myelinated fibers, which mandates that the regression slope < 1 (i.e. $D_F > D_A$), excludes regression fits for $\lambda > 1$.

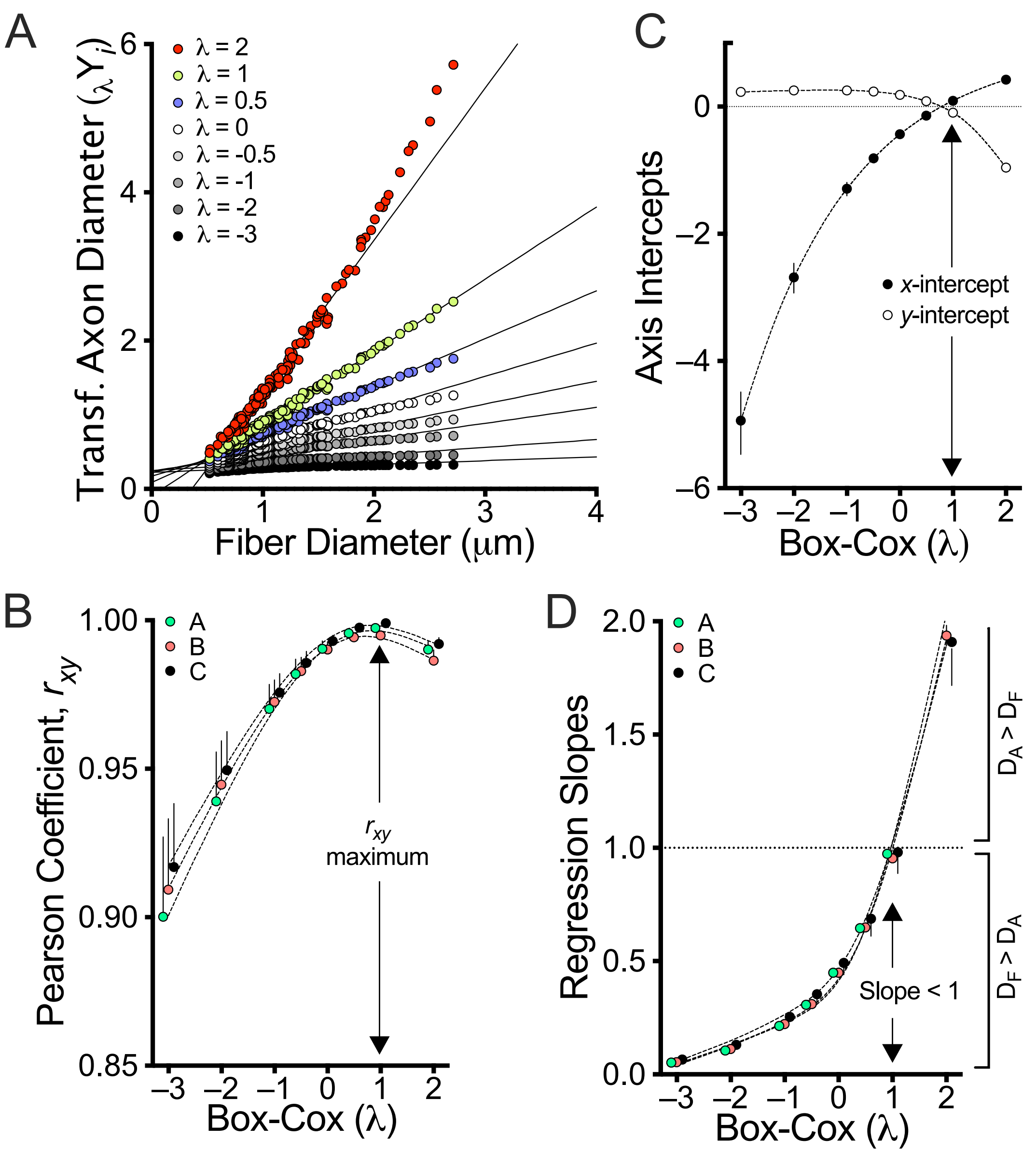


*Supplemental Figure 3 – statistical and biological constraints on rsh axon-fiber diameter plots*
**A.** Summary of a Box-Cox lambda transformation series for axon diameters from mouse D plotted against the respective fiber diameters (Fig. 1B). Each transformation is fit using linear regression to estimate parameters that identify the most likely axon-fiber diameter relation (linear versus nonlinear). **B.** Pearson's correlation coefficients for the regression fits (mice D-F) are plotted against the corresponding Box-Cox lambda values. The maximum $r_{xy}$ value, for $\lambda = 1$, is the most likely axon-fiber diameter relation as confirmed by mixed-effects analysis with Geisser-Greenhouse correction ($F_{(1.049,2.098)} = 60$, $\varepsilon = .15$, $P = .014$). Post hoc Sidak's multiple comparison's tests indicates that $r_{xy}$ for $\lambda = 1$ is only statistically different from $r_{xy}$ for $\lambda = -3$ ($P = .043$). But more importantly, no other values of lambda yield superior correlations, which supports the null hypothesis that $\lambda = 1$ is the most likely regression fit. **C.** The *y*-intercepts ±95% CI for the regression fits in (A) plotted against the Box-Cox lambda values. Biological constraints on the regression fits – *x*-intercept $\leq 0$ and *y*-intercept $> 0$ – exclude transformations of $D_A$ where $\lambda < 1$. **D.** The constraint of myelinated fibers, which mandates that the regression slope < 1 (i.e. $D_F > D_A$), excludes regression fits for $\lambda > 1$.

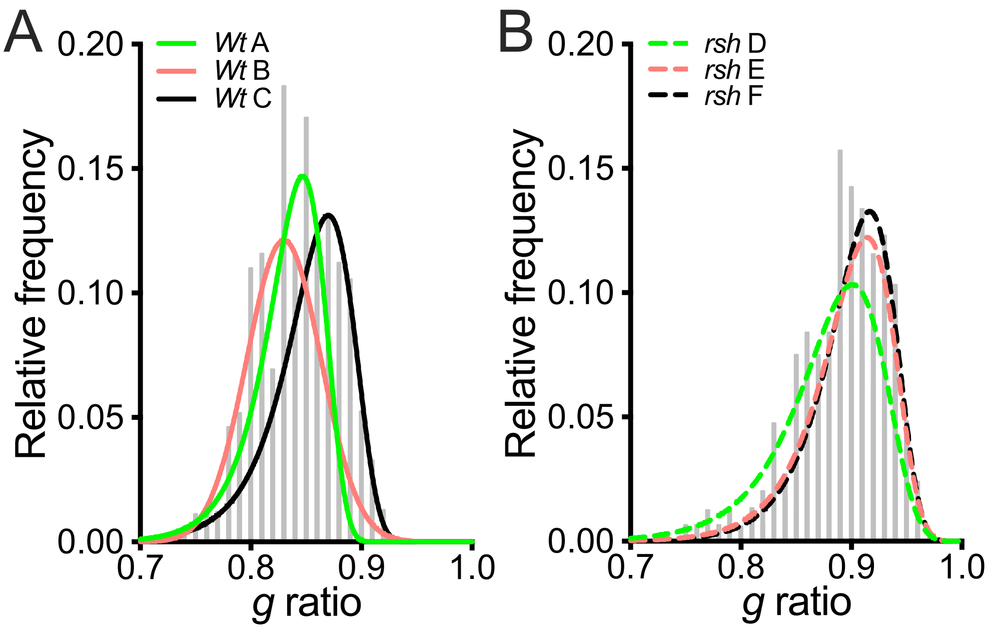


*Supplemental Fig. S4 – g ratio relative frequency histograms reveal skewed data structure*
Despite *g* ratios being superseded as an estimator for the axon-fiber diameter relation, there is reason to persist with this metric in future studies; not for the traditional purpose, but rather as a qualitative estimator for skewness. Indeed, these ratio data provide important general insights into data structure[4]. **A.** Figs 1A and S2 demonstrate that the axon-fiber diameter relation in wild type optic nerve is directly proportional, and a relative frequency histogram of the corresponding *g* ratios is expected to be roughly symmetric, which is the case (Gaussian fits). **B.** The *rsh* mouse data are conspicuously non-Gaussian (Gumbel distribution fit). This left skewing does not directly reflect the distributions for the axon, fiber or myelin diameters, all of which are actually right skewed (Fig. S1). Nevertheless, several properties of the axon-fiber relation contribute to *g* ratio left skewing. For example: all *g* ratios are between 0.7-1.0 which in general compresses the right tail of the distribution because the ratio transformation is nonlinear (called the boundary effect[4], *g* ratios < unity by definition); hypomyelination exacerbates the proportion of *g* ratios near unity; the loss of direct proportionality in the axon-fiber diameter relation (Fig. 1F). While important for detecting experimental artifacts that cause *g* ratios to appear correlated with fiber diameter[2,6], relative frequency histograms or other skewness metrics like Pearson's $\gamma_1$, can be unreliable[7]. Thus, a better approach is to avoid data skewing by computing $g_c$ ratios or using the *g'* cline.

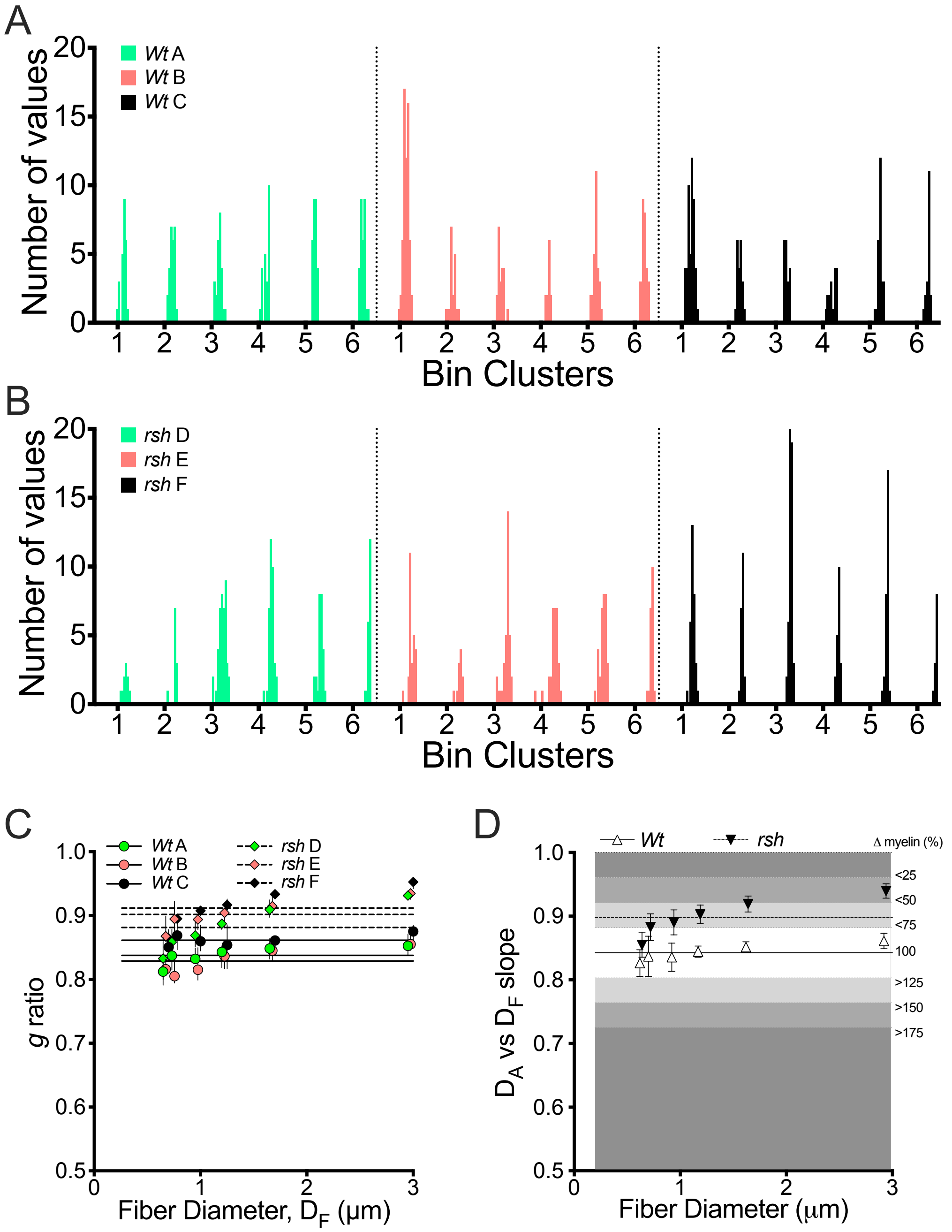


*Supplemental Figure 5 – g ratio pipeline analysis for the current study data*

**A,B.** Histograms from wild type mice A-C (A) and *rsh* mice D-F (B) show that *g* ratios in most bin clusters are unimodal with normal or skewed distributions which can be summarized as medians ±95% CI for each mouse. Most clusters have roughly equal numbers of *g* ratio values. **C.** median *g* ratio values ±95% CI for the wild type and *rsh* mice and the corresponding horizontal regression fits. While most of the these fits are within the 95% CI limits for the wild type mice (i.e. *g* ratios are independent of fiber diameter), this does not appear to be the case for the *rsh* mice. **D.** Summary of the data in panel (C), which is the final output figure from the *g* ratio analysis pipeline[6]. Compared to the wild type cohort, the *rsh* mice have approximately 60% less myelin, which largely comports with Fig. 4 and the previous pipeline study[6]; however, the current study demonstrates that the apparent slope in the *rsh* data is an artifact stemming from a loss of direct proportionality in the *rsh* axon-fiber diameter relation (Fig. 1). On the other hand, the *g'* cline for this cohort reflects the linearity of the axon-fiber relation (Fig. 3).

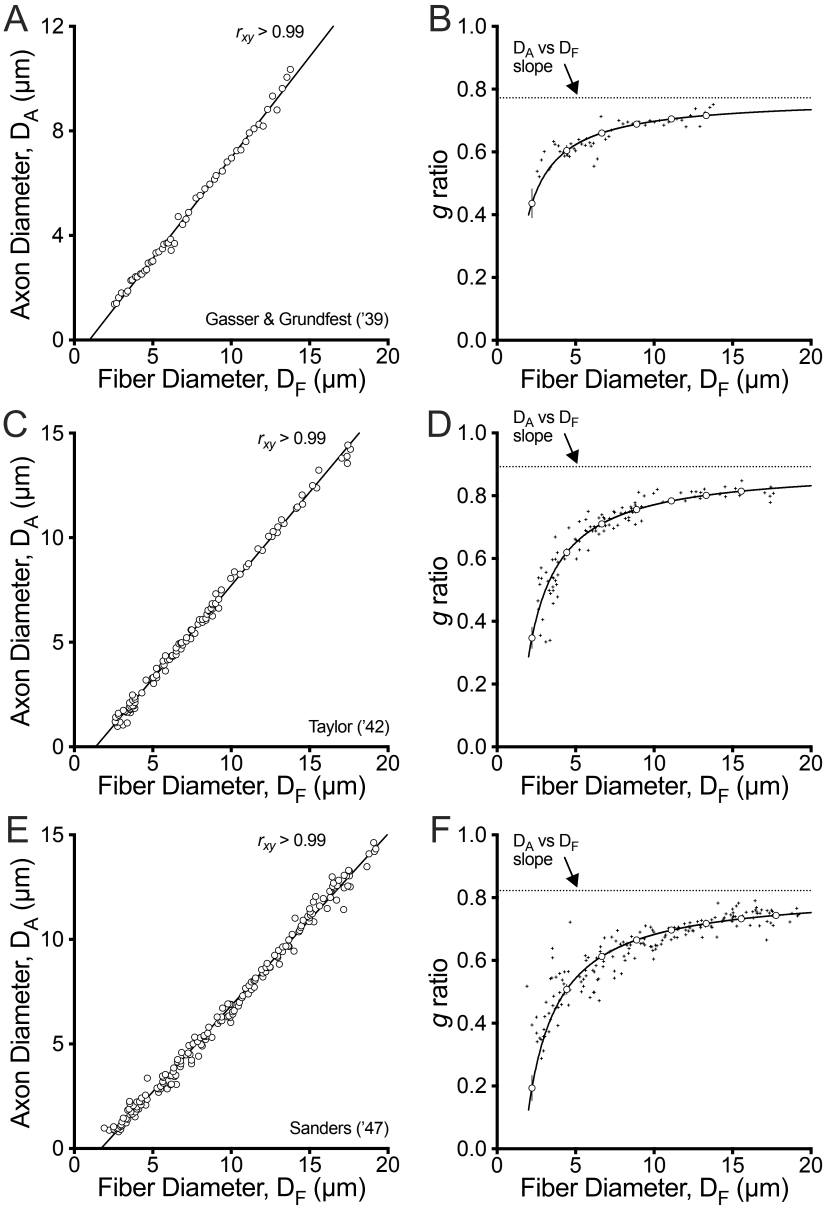


*Supplemental Figure 6 – pioneering studies of PNS myelin g ratios in vertebrates*
Scatterplots from three early studies[1,8,9] were digitized using WebPlotDigitizer ver4.7 (https://automeris.io/WebPlotDigitizer). Axon, fiber and myelin diameters as well as *g* ratios were computed. The digitized data indicate that significant artifacts were present. For example, the studies by Taylor and Sanders included PNS myelinated axons down to 0.2μm diameter, an obvious systematic error caused by insufficient measurement resolution under light microscopy (PNS axons significantly below 1 μm diameter are unmyelinated[3]). To minimize the effects of this artifact, axons < 0.8μm diameter were excluded from the digitized data prior to plotting. Unfortunately, such technical errors have played a significant role in misunderstanding and misinterpretation of *g* ratio plots for decades. **A,C,E**. Because of non-zero *y*-intercepts in these plots, the corresponding *g* ratio plots **B,D,E**, are curvilinear, and are fit using a reciprocal function (eqn 5, Methods). Per Fig. 2, curvilinearity reflects *g* ratio underestimates of the regression slope in the axon-fiber diameter scatterplots. (A,B) Data (675 fibers summarized into 52 equally-sized bins in the original study) for saphenous nerve of cat from figure 12 in Gasser & Grundfest[1]. (C,D) Data (113 fibers) for saphenous and sciatic nerves of the cat from figure 3 in Taylor[8]. (E,F) Data (200 fibers) for peroneal nerve of the rabbit from figure 3 in Sanders[9].

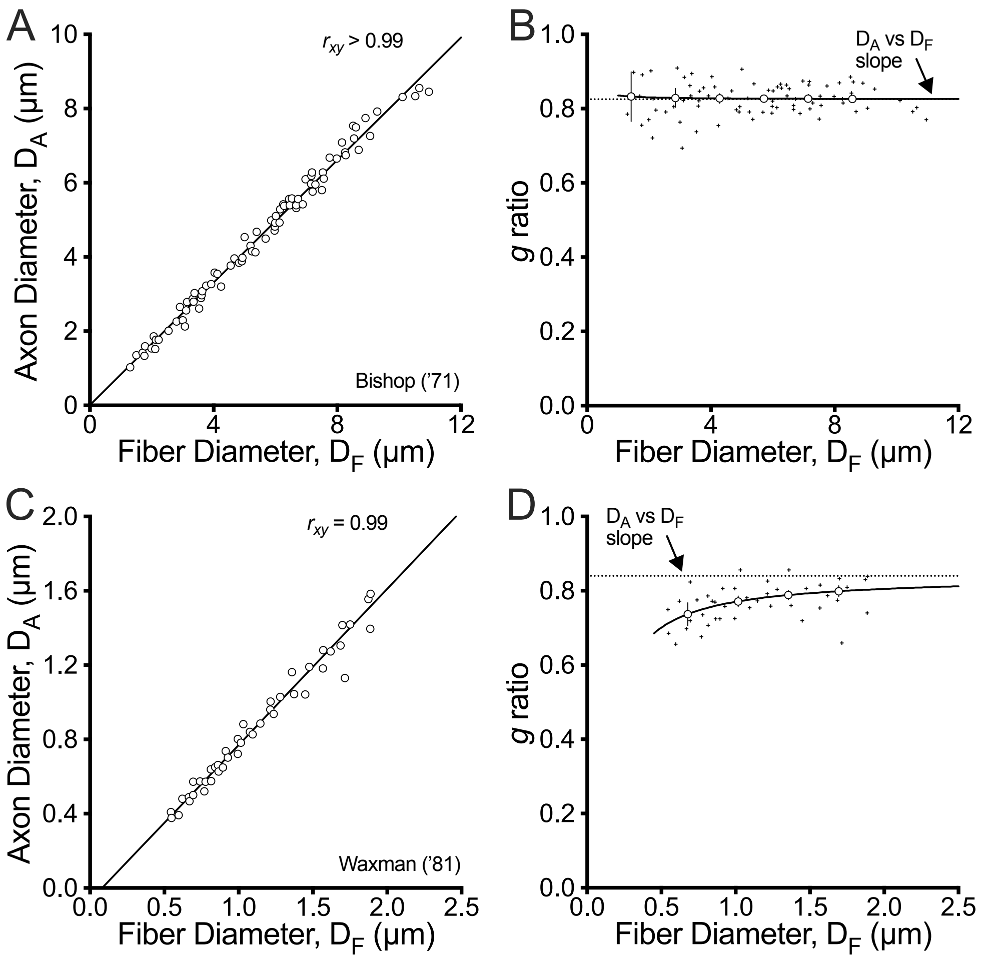


*Supplemental Figure 7 –pioneering studies of CNS myelin g ratios in vertebrates*
Scatterplots from two electron microscopy based studies[10,11] were digitized using WebPlotDigitizer ver4.7 (https://automeris.io/WebPlotDigitizer). Axon, fiber and myelin diameters as well as *g* ratios were computed. Axon, fiber and myelin diameters as well as *g* ratios were computed. The scatterplots from these studies are relatively artifact-free. **A,B.** Data (81 fibers) for the fasciculus gracilis of the cervical dorsal funiculus in cat from figure 1 of Bishop and colleagues[10].The regression fit to the axon-fiber diameter scatterplot (A) passes through the Origin and the corresponding *g* ratio plot (B) has essentially zero slope in accordance with the axomyelin unit model. **C,D.** Data (45 fibers) for the splenium of corpus callosum in rhesus monkey from figure 2 of Waxman[11] has a small y-intercept offset.

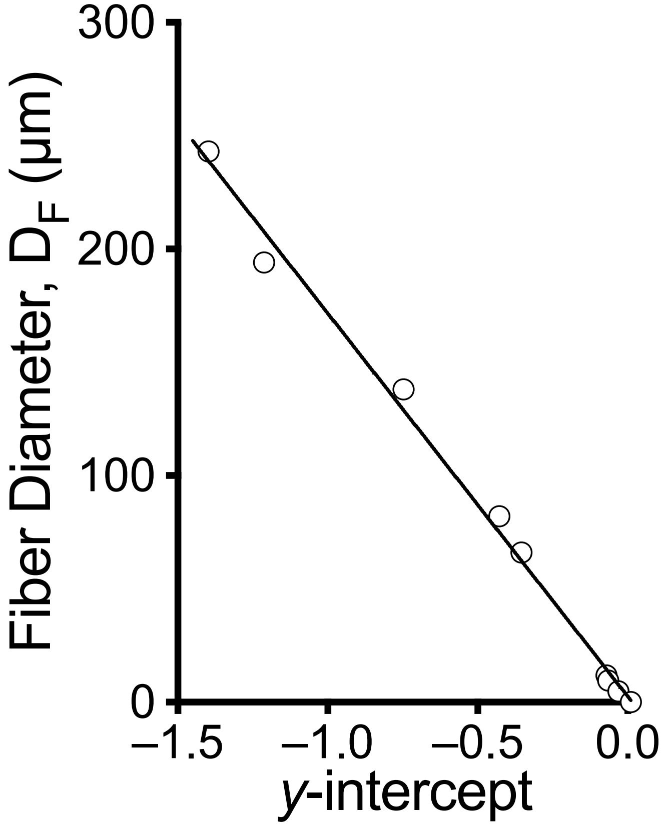


*Supplemental Figure 8 – the magnitude of the y-intercept determines the caliber of fibers above which g ratios become constant*

The notion that *g* ratios are invariant for fibers larger than 6-10μm, while they decrease progressively for smaller fiber calibers (i.e. myelin sheaths are disproportionally thick for small fibers), is controversial and has been thoroughly reviewed and discounted[5]. The current study also discounts this notion, and demonstrates why it lacks merit. Thus, the results in Table S1 were computed from Figs 1A, 1B, 2A, 2B, S6 and S7 using eqns S12-14 in Supplement S7, and plotted to demonstrate the inversely proportional ($r_{xy}$ < 0.99) relation between *y*-intercept and the caliber of fibers above which *g* ratios are invariant. The threshold fiber diameter for this invariance was determined by extrapolating the reciprocal function regression fit for the *g* ratio plots, and was defined as a value within 1% of the *g'* cline.

Table S1 – estimates of the minimum fiber diameter at which *g* ratios become constant (i.e. within $\delta = 0.7\%$ of the axon-fiber diameter slope)

| Published data or figure in the current study | $m_a (= m_b)$ | $(1-\delta) * m_a$ (eqn S18) | $y\text{intercept}_b$ (eqn 5) | $D_F$ (μm) (eqn S19) |
|---|---|---|---|---|
| Schmitt & Bear[12] | 0.748 | 0.743 | -0.427 | 82 |
| Gasser & Grundfest[1] | 0.772 | 0.767 | -0.746 | 138 |
| Taylor '42[8] | 0.893 | 0.886 | -1.211 | 194 |
| Sanders[9] | 0.822 | 0.817 | -1.397 | 243 |
| Bishop et al[10] | 0.825 | 0.819 | 0.011 | 0 |
| Waxman[11] | 0.840 | 0.834 | -0.070 | 11.8 |
| Graf von Keyserlingk & Schramm[13] | 0.768 | 0.763 | -0.353 | 66 |
| Wt A-C, Figs 1A, 2A | 0.873 | 0.867 | -0.031 | 4.92 |
| *rsh* D-F, Figs 1B, 2B | 0.966 | 0.951 | -0.065 | 9.68 |
